# A keratin bundling transition uncages the nucleus in stretched epithelia

**DOI:** 10.1101/2025.08.25.672104

**Authors:** Tom Golde, Marco Pensalfini, Nimesh Chahare, Pere Roca-Cusachs, Gerhard Wiche, Guillaume T. Charras, Marino Arroyo, Xavier Trepat

## Abstract

There is broad consensus that intermediate filaments play a key role in protecting cells and tissues from large deformations. However, little is known about how they fulfil this function. Here we demonstrate that epithelial cells undergo a slow adaptation to stretch that couples a star- bundling transition of keratin filaments and the escape of the nucleus from its keratin cage. The bundling transition begins with a depletion of keratin filaments at tri-cellular junctions followed by a progressive accumulation in thick bundles that bisect cell-cell junctions. Bundling is a cooperative process that initiates in a few scattered cells and propagates to their neighbours through desmosomes, leading to the growth of multicellular clusters that contain a percolated network of thick keratin bundles. Bundling dynamics are slow and strongly influenced by the interaction between actin and keratin. Informed by a computational model, we provide evidence that keratin bundling generates a compressive stress on the nucleus, which is relaxed by nuclear escape from the keratin cage. The topological transitions identified here provide epithelia with a multiscale mechanism to adapt to sustained stretch.

## Introduction

Intermediate filaments, such as keratin and vimentin, constitute an essential class of cytoskeletal polymers widely expressed in animal cells. Owing to their hierarchical structure, individual filaments and their networks exhibit exceptional mechanical properties: they can stretch several times their original length but undergo pronounced stiffening at higher strains^1–5^. At the single cell level, intermediate filaments support cell integrity by resisting both tensile and compressive stresses and shielding the nucleus from its mechanical microenvironment ^6–12^. In epithelial tissues, keratin intermediate filament networks are responsible for strain-stiffening and crucial to maintaining tissue cohesion during large deformations ^13–17^. Collectively, these findings have led to the growing consensus that, while actin filaments largely determine cell and tissue mechanics at small deformations, intermediate filaments play a key role in governing cellular responses to larger strains ^17–20^.

While extensive functional and mechanical evidence now support this consensus view, the reorganization of intermediate filament networks in response to mechanical stretch and their specific role in providing mechanical protection remain poorly understood. In relaxed epithelial monolayers, keratin filaments form two interconnected networks, one lining the cell cortex and another enveloping the nucleus. The cortical keratin network, commonly referred to as the rim, connects adjacent cells through desmosomes and is linked to the nuclear-enveloping network through short filament bundles known as spokes. This rim-and-spoke structure relies heavily on the crosslinking protein plectin, which bridges the actin and keratin cytoskeletons and couples keratin to the nucleus via nesprins ^21,22^. Upon mechanical stretching, the organization of these networks is transformed ^23–28^. In MDCK monolayers, cells exposed to large strains display long, thick keratin cables that withstand tension and form supracellular networks ^19,29^. Long keratin cables spanning several cells have also been found in developing embryos, suggesting a protective and stabilizing role ^30^. How such keratin structures form under stretch, how they organize into supracellular networks, and how they modulate the mechanical properties of the tissue is not known.

Here we combine experimental and theoretical approaches to study the dynamic reorganization of the keratin cytoskeleton and its coupling to the nucleus during sustained stretching of epithelial monolayers. Using a new device to stretch curved cell monolayers, we found that the keratin network adapts to prolonged mechanical loading by undergoing a slow bundling transition. The kinetics of this transition are largely governed by the actin-keratin interaction. Strikingly, we show that the bundling transition leads to the full escape of the nucleus from its keratin cage. A computational model explains this behaviour in terms of a compression-by-tension mechanism in the entangled keratin network. Finally, we demonstrate that the drastic structural remodelling of the keratin cytoskeleton is a collective supracellular process driven by a nucleation-and-growth mechanism. Our study unveils a new multiscale response of cellular monolayers to sustained stretch, wherein the nucleus is ultimately unshielded from the keratin cytoskeleton.

## Results

### Keratin undergoes star-bundling transition in spontaneous domes

We first studied epithelial domes that form spontaneously in MDCK monolayers due to the osmotic pressure generated by directional ion pumping ^31,32^. These domes undergo autonomous cycles of inflation and deflation, lasting several hours and reaching areal strains of 200-300%. Using micropatterning techniques, we generated arrays of spontaneous domes with a defined footprint radius^19^ and fixed them to visualize the spatial distribution of F-actin and keratin-18 filaments (Fig. 1a, Supplementary Fig.1a). While the distribution of F-actin was largely cortical in all cells, keratin displayed two markedly distinct organizations depending on the extent of stretch. Cells within the unstretched monolayer showed a prominent keratin mesh enveloping the nucleus, connected to the cell surface through thin and short bundles (Fig. 1b, Supplementary Fig. 2). These unstretched cells also displayed a thin keratin network lining the cell cortex, consistent with the rim-and-spoke arrangement commonly reported in epithelial cells ^33,34^. In contrast, stretched cells within the dome showed a star-like arrangement, composed of a dense central node and long, thick bundles radiating towards the cell periphery (Fig. 1b). Each bundle was mirrored by a corresponding bundle in the neighbouring cell, leading to the emergence of a supracellular percolated network of interconnected bundles. This network defines supracellular clusters bounded by transitional cells which exhibited partial bundling restricted to specific subcellular regions.

**Fig. 1:**
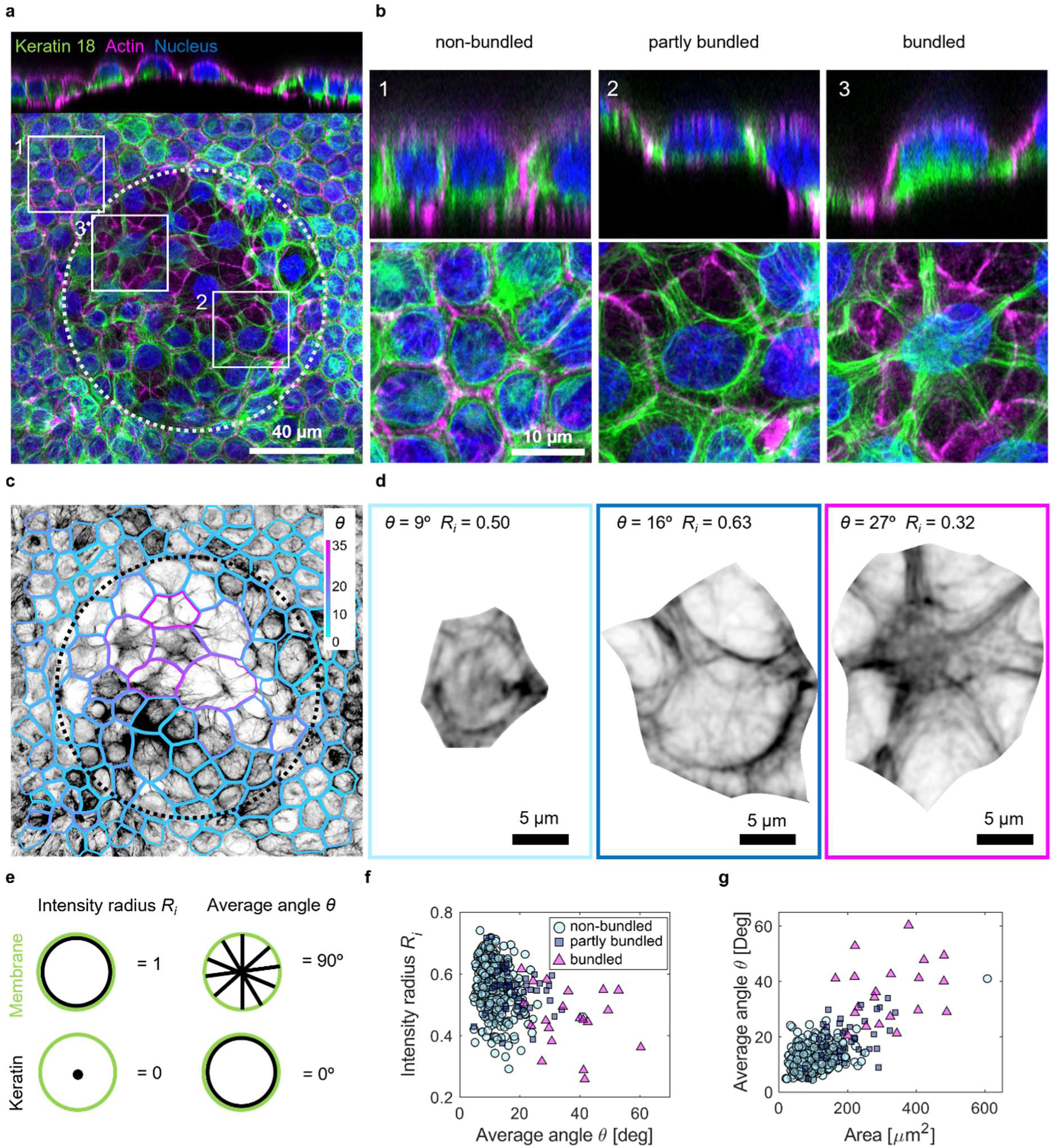
K**e**ratin **bundles in spontaneous domes. a,** Spontaneous dome of MDCK cells expressing keratin-18-GFP (green) stained for F-actin (magenta) and nuclei (Hoechst, blue). Top: lateral view. Bottom: maximum intensity projection. **b,** Zoomed views of the three regions marked with white rectangles in **a**, showing non-bundled, partially bundled, and bundled cells. Top: lateral view. Bottom: maximum intensity projection. **c,** Keratin-18-GFP intensity in the spontaneous dome shown in **a**. The segmented cell boundaries are color-coded according to the value of the average angle *θ*. **d,** Keratin-18-GFP image of the three cells shown in **b**. **e,** Schematic illustration of the intensity radius *Ri* and average angle *θ*. **f**, Intensity radius plotted against the average angle (R=-0.34, p=2.9E-15). **g**, Average angle *θ* plotted against cell area (R=0.71,p=6.3E-79). n=509 cells in **f** and **g**.

To assess the bundling state of each cell from keratin images, we defined two metrics (Fig. 1e). One is the intensity radius (*Ri*), reflecting the radial distribution of keratin within the cell. Values close to 1 indicate a predominantly peripheral localization, while values near 0 indicate a central localization. The second metric is the average angle (*θ*), quantifying orientation relative to the cell tangent for keratin filaments located in the outer half of the cell. This metric approaches 90° for filaments organized radially and 0° for those arranged circumferentially (Fig, 1e, Supplementary Fig. 3). Non-bundled cells displayed high values of *Ri* and low values of *θ*, highlighting that filaments accumulate close to the cell junctions and arrange parallel to it. Conversely, cells forming star-like bundles showed low values of *Ri* and high values of *θ*, reflecting the central accumulation of keratin and the radial orientation of the bundles. Partially bundled cells displayed intermediate values of *θ* and values of *Ri* similar to non-bundled cells, reflecting the absence of a central knot (Fig. 1c-f). The analysis of these metrics across all cells in the domes established a clear dependence between cell area and its bundling state. Indeed, by plotting *θ* vs cell area we found that small cells are generally non-bundled, large cells are bundled, and intermediate-size cells are partially bundled (Fig. 1g). This result prompted us to study the mechanisms underlying bundle formation and their dynamics.

### Dynamics of the star-bundling transition

In previous theoretical work, we hypothesized that bundling could be explained as a topological phenomenon in an entangled network of filaments connected to laterally mobile desmosomes, where the straightening of keratin filaments upon stretch is ultimately hindered by a self-organized knot ^35^. To test this hypothesis, we developed a microfluidic device to form domes with defined geometry and dynamic control of inflation (Fig. 2a, Supplementary Fig. 1b). This device consists of two perpendicular channels separated by a thin PDMS layer containing evenly spaced holes with a diameter of 80 µm that define the dome footprint. This PDMS layer is covered with a permeable polycarbonate membrane featuring 400nm-sized pores that allow the passage of liquid but not of cells. Using a photopatterning technique (PRIMO^36^), the polycarbonate membrane is coated with a high concentration of fibronectin outside the footprint and a low concentration of fibronectin inside of it. This enables the seeding of cells and the subsequent formation of a confluent monolayer on the fibronectin-coated side. Hydrostatic pressure can then be applied by injecting fluid beneath the membrane, causing a delamination of the monolayer only at the low adhesion footprint. With this approach, we can form directed epithelial domes of defined geometry and control their luminal pressure over time (Fig. 2a, Supplementary Fig. 1b).

**Fig 2:**
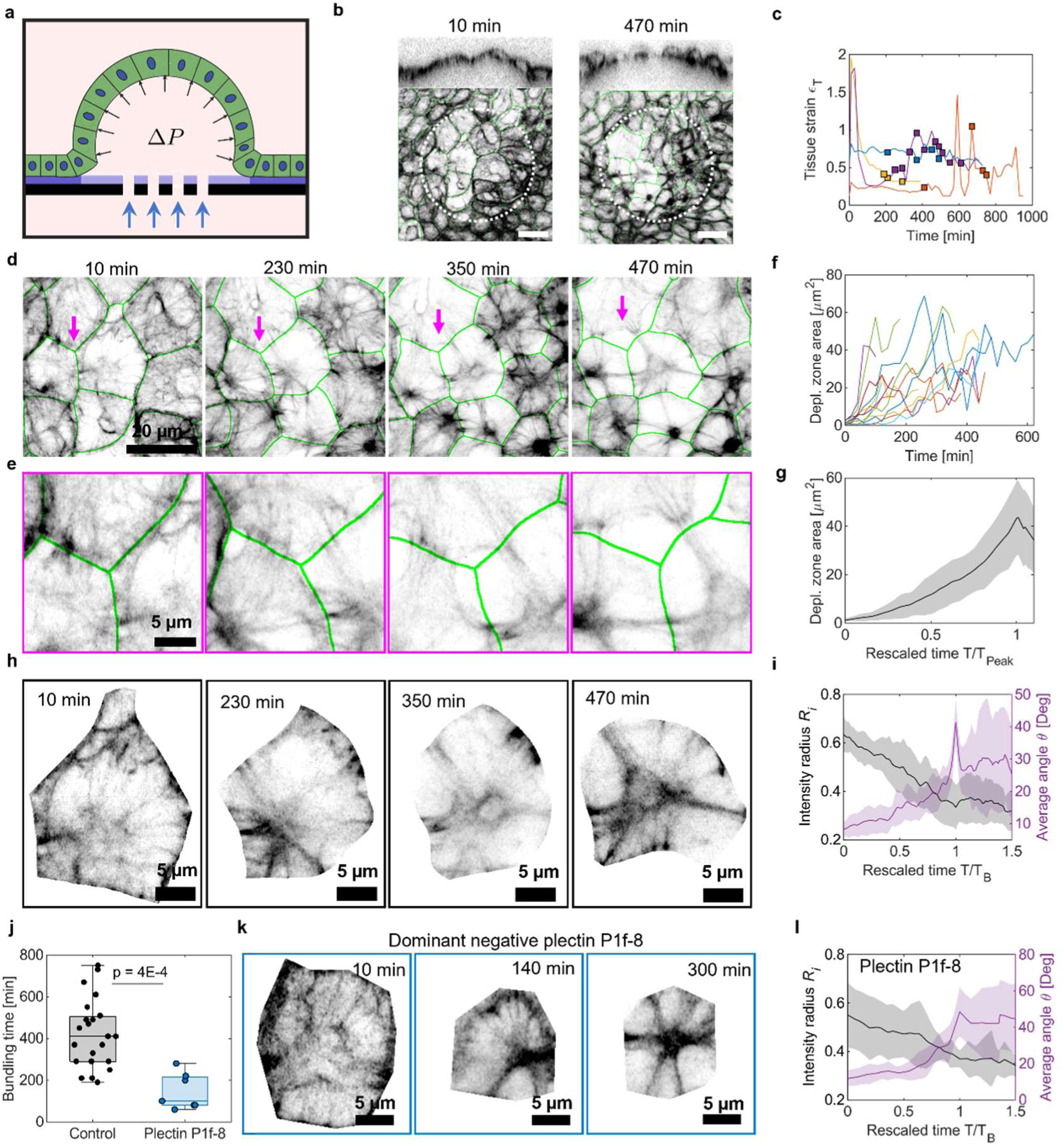
B**u**ndling **dynamics in directed domes. a,** Scheme of the microfluidic device showing the porous membrane (black) coated with high (dark violet) and low (light violet, footprint) fibronectin levels. Hydrostatic pressure applied from the basal side causes the delamination of the cell monolayer at the footprint, resulting in the formation of a dome. **b,** Maximum intensity projection and side view of a directed dome of keratin-18-GFP cells. The dome footprint is marked with a dotted white circle and segmented cell borders are marked in green. **c**, Time evolution of tissue strain for 4 independent domes. Squares mark each bundling event, defined as an individual cell crossing the bundling threshold. **d,** Time sequence of cropped images of a directed dome. Arrows indicate the formation of a keratin depletion zone at a tricellular junction. **e,** Zoomed view of the tricellular junction marked in **d**. **f**, Time evolution of the depletion zone area. **g**, Mean depletion zone area as a function of time rescaled by the peak time (n = 13 cells). **h**, A representative individual cell undergoing the bundling transition. **i**, Mean intensity radius *Ri* (grey) and mean average angle *θ* (violet) plotted against time rescaled by the bundling time (n = 23 cells). **j**, Bundling time for control cells and cells overexpressing plectin P1f-8-BFP. Boxes indicate the median and interquartile range, and whiskers extend to minimum and maximum values. **k**, A representative individual cell overexpressing plectin P1f- 8-BFP undergoing the bundling transition. **l**, Same as (**i**) but for P1f-8-BFP cells (n = 7 cells). Shaded areas in (**g,i,l**) represent the standard deviation.

Using this device, we applied a constant pressure of 200 Pa and performed confocal timelapse imaging of keratin-18-GFP and a cell membrane marker for several hours (Fig. 2b). Despite the application of constant pressure, dome strain fluctuated substantially over time, likely due to discrete cell division and extrusion events as well as transient paracellular leaks. (Fig. 2c). Contrary to our expectations, keratin bundling did not occur immediately after stretch application (Fig. 2b,d and Supplementary Video 1). Instead, only after tens of minutes, we observed the slow formation of keratin-depleted zones at tri-cellular junctions (Fig. 2d,e and Supplementary Video 2). These zones gradually grew over time, reaching an average area of 43 ± 15 µm^2^ (mean ± s.d.) within 4.5 ± 2.3 h (mean ± s.d.). Keratin filaments depleted from these zones progressively moved towards the perpendicular bisector of each cell-cell junction, ultimately leading to the star-shaped geometry characteristic of stretched cells (Fig. 2 d-g). While keratin depleted zones appeared as a gap in the monolayer, the actin cortex and cell-cell junctions remained continuous, indicating that epithelial integrity was preserved (Fig. 1b). We thus conclude that tricellular junctions are the nucleation sites of a structural reorganization of the keratin cytoskeleton, which directly initiates the bundling transition.

To quantify the dynamics of keratin bundling we computed the time evolution of *θ* and *Ri*, and defined the bundling transition using a threshold based on these two metrics (see methods for details). The average bundling time was 424 ± 162 min (mean ± s.d.), with substantial variability between cells (Fig. 2j). To analyse systematically the dynamics of bundle formation across the cell population, we plotted *θ* and *Ri* as a function of time rescaled between 0 and 1, where 0 corresponds to the onset of pressure application and 1 to the completion of the bundling transition. Cells undergoing the bundle transition showed a gradual decrease of *Ri*, from predominantly peripheral (>0.5) to central (<0.5), while *θ* increased smoothly. Notably, *θ* showed a sharp rise shortly before crossing the bundling threshold (Fig 2h,i), revealing accelerated cytoskeletal dynamics just before the transition. After crossing the threshold, *θ* remained high and *Ri* remained low, showing that cells retained the bundling state. Even after spontaneous dome deflation and reinflation episodes, bundling was not lost (Supplementary Fig. 4), indicating that the bundling transition is irreversible within the time scales of our study.

In summary, our findings indicate that the star-bundling transition is not an immediate reaction to the increase in cell area, but rather a slow adaptation to stretch over hundreds of minutes. Therefore, the previously reported strain stiffening behaviour provided by keratin in epithelial cells at short time-scales (<1min) ^17^ is independent of the bundling transition.

### Bundling is hindered by the actin cortex

We next investigated the mechanisms that explain the slow dynamics of the bundling transition. An appealing candidate is the interaction between the intermediate filaments and the actin cytoskeleton, which is well-known to influence cytoskeleton architecture ^37–39^. To study the role of this interaction, we targeted plectin, the main crosslinking protein between keratin and actin^22,40,41^. Specifically, we induced a loss of plectin function by overexpressing a truncated version of the plectin isoform P1f tagged with BFP (P1f-8-BFP). This mutant retains the actin-binding domain but lacks the keratin-binding domain, thereby acting as a dominant negative mutant that weakens the specific interaction between the actin cortex and the keratin cytoskeleton ^38^. In MDCK cells, P1f-8-BFP localizes mainly at the cell cortex (Supplementary Fig. 5).

In the absence of stretching, cells expressing P1f-8-BFP and control cells exhibited comparable keratin organization (Supplementary Fig. 5). Upon stretching, the P1f-8-BFP cells underwent a bundling transition similar to that observed in control cells (Fig. 2 k,l). However, the average bundling time was significantly reduced to 146 ± 68 min (mean ± s.d.) (Fig. 2j). This near 3-fold decrease establishes that the keratin-actin interaction is a main determinant of the bundling rate. To further investigate the role of the actin cortex during the bundling transition, we treated control MDCK domes with 1µM Latrunculin A 15 min after dome formation, causing a rapid disassembly of actin filaments. This perturbation resulted in a quick reorganization of the keratin cytoskeleton into a more bundled network in less than 60 min, although the characteristic star-like bundle organization observed in untreated cells was not fully recapitulated (Supplementary Fig. 6). These experiments establish that actin-keratin interactions are a major determinant of the bundling transition during stretch.

### A computational model captures the bundling transition and predicts nuclear uncaging

Aiming to recapitulate the dynamics of the bundling transition, we expanded our previous computational framework, which was constrained to a 2D geometry and did not account for keratin interactions with the actin cytoskeleton or the nucleus ^35^. We developed a customized version of the open-source simulation suite Cytosim^42^, which enables stochastic simulations of cytoskeletal fibres governed by Brownian dynamics (Supplementary Note). Our new model simulates a representative cell comprising flexible and entangled keratin fibres whose ends are attached to the cell periphery via laterally mobile bonds representing desmosomes. Fibres interact with each other through steric repulsion forces, which prevent filament crossing, and short-range adhesive forces, which promote bundling.

Within this computational framework, we first implemented a 2.5D simulation of the region around the medial plane of the cell (Fig. 3a, Supplemental Fig. 7a,b), where the rim-and-spoke organization is better observed experimentally. In this 2.5D model, the nucleus is represented by a cylindrical region that cannot be penetrated by keratin fibres. Keratin interactions with the nuclear surface and the cell cortex are modelled by uniformly distributing a large number of discrete stochastic linkers on both surfaces. These linkers represent plectin proteins that connect keratin with the actin cortex by binding to actin filaments and with the nuclear envelope by binding to nesprin-3^43^. In the model, the linkers bind fibres stochastically with binding rate 𝐾^+^ and unbind as slip bonds with a force-dependent rate. As the nucleus moves or the cellular surfaces deform, the position of these linkers evolves accordingly. Remarkably, the introduction of such linkers into an initially disordered keratin network was sufficient to drive its spontaneous reorganization into a rim-and-spoke architecture after an equilibration period 𝑡_eq_ ≫ 1/𝐾^+^ (Supplementary Fig. 7c). The emergence of such a characteristic arrangement was robust across a range of parameter choices, suggesting that the rim-and-spoke topology is self-organized and arises generically from interactions between keratin fibres and the intercellular and nuclear ‘sticky’ surfaces bounding the network.

**Fig 3:**
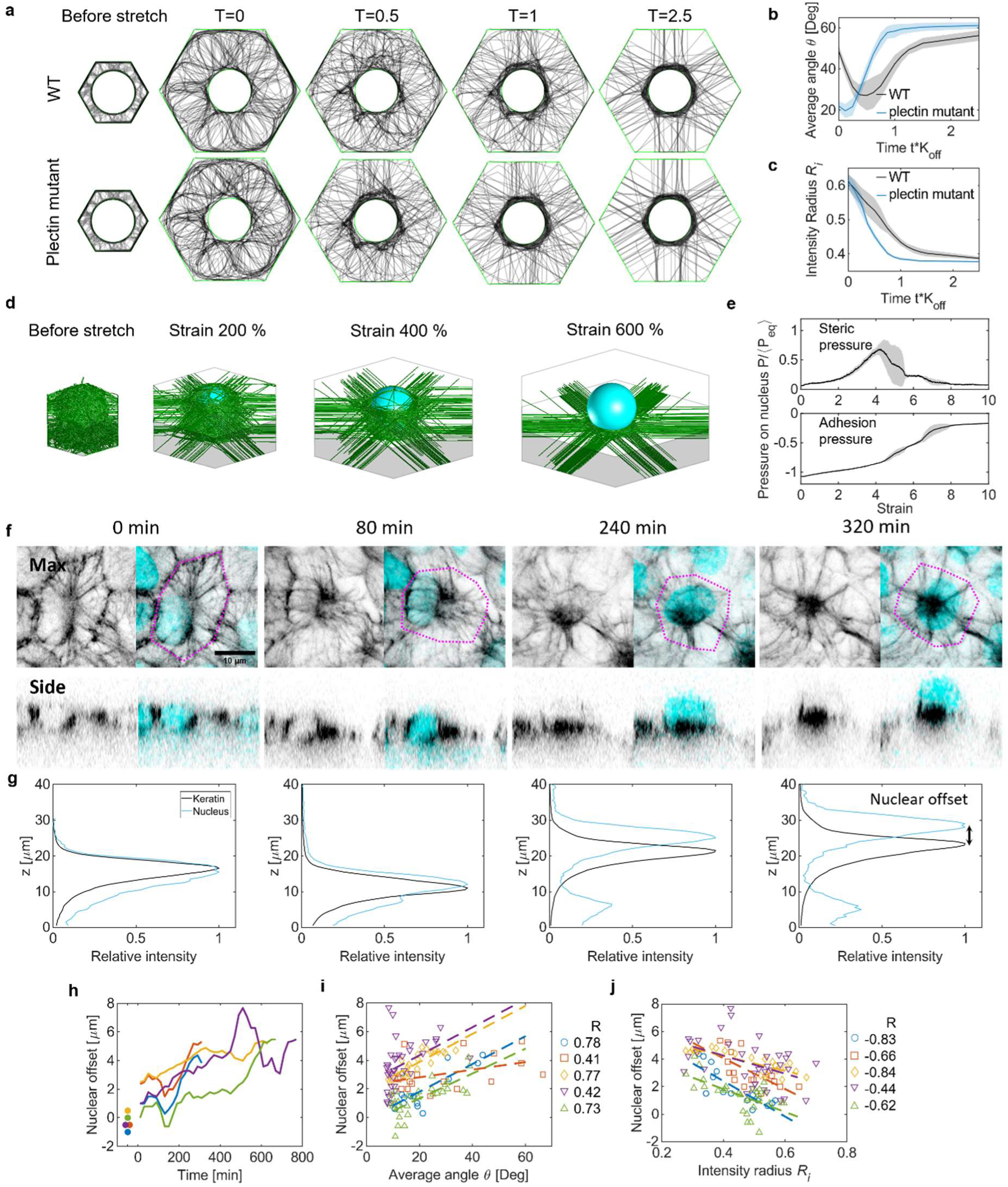
T**h**e **nucleus escapes from the keratin cage during bundling. a,** Top view of a 2.5D simulation with keratin filaments represented in black and cell boundaries marked in green**. b,** Mean simulated average angle *θ* plotted against time, reported as mean ± s.d. of n=8 model realizations. **c**, Mean simulated intensity radius *Ri* plotted against time, reported as mean ± s.d. of n=8 model realizations. **d**, Isometric view of a 3D simulation with keratin (green) and nucleus (cyan). **e**, Mean simulated pressure on the nucleus plotted against strain in the 3D simulation, reported as mean ± s.d. of n=7 model realizations and with pressure values normalized with respect to the time average of their sum during the equilibration phase preceding cell stretching. **f,** Maximum intensity projection and side view of a MDCK keratin-18- GFP cell (grey) showing the nucleus (SPY-DNA, cyan) and cell border (magenta dotted line). **g**, Normalized intensity of the keratin and nuclear signals along the z coordinate. **h**, Nuclear offset plotted against time. Dots indicate the values in the relaxed monolayer before dome inflation. Different colours represent different cells. **i**, Nuclear offset plotted against the average angle *θ*. **j**, Nuclear offset plotted against the intensity radius *Ri*. Dashed lines in (**i,j**) indicate a linear fit. (n = 5 cells in **h,i,j**). R values are the Pearson correlation coefficient.

Next, we subjected the model to in-plane loading by applying a stretch and hold protocol. In the control case, stretching led to a gradual loss of the rim, the formation of load bearing radial bundles, and an accumulation of fibres around the nucleus, which we quantified plotting *θ* and *Ri* as in the experiments (Fig. 3a-c). To test the effect of the plectin loss of function, we reduced the number of discrete linkers by 50%. In this case, stretching led to the same characteristic reorganization of the keratin structure but on an accelerated time scale (Fig. 3a-c). This confirms that keratin-actin interactions play a key role in modulating the timescale of the bundling transition. However, the simulation never reached the full star-like organization of the keratin network due to the geometric obstacle posed by the nucleus in the 2.5D model.

To address this limitation and better investigate the interaction between keratin and the nucleus, we developed a second set of 3D models. In these models, the cell is a hexagonal prism, and the nucleus is a rigid sphere with translational degrees of freedom. The nucleus interacts sterically with entangled keratin fibres attached to the side surfaces of the cell via laterally mobile links (Supplemental Fig. 7d). To account for the apical-basal asymmetry of epithelial cells, we introduced a confining potential for the nucleus on the bottom and side walls of the cell membrane but not on its apical surface. Given the computational complexity of the 3D models, we simplified interactions between keratins and other cytoskeletal components by introducing short-range adhesive potentials on the nuclear shell and on the bottom and side cell surfaces. We first confirmed that this modelling approach also captures the rim-and-spoke keratin arrangement (Supplemental Fig. 7d). We then gradually stretched the model at a constant strain rate. As strain increased, keratin reorganized into a star-like bundle structure and keratin fibres apically covering the nucleus were progressively lost. This was followed by a rapid escape of the nucleus from the keratin cage as the perinuclear network slid basally to form the central knot (Fig. 3d and Supplementary Video 3).

We then wondered about the physical forces exerted by the keratin fibres on the nucleus during this uncaging process. To this end, we computed two contributions to nuclear pressure: the adhesion pressure, which results from the adhesive interaction between the nucleus and the keratin network, and the steric pressure, which arises from the fact that keratin fibres are excluded from the nucleus. Before stretch, the steric pressure was nearly zero, whereas the adhesion pressure was negative, reflecting the tension of the ‘spoke’ fibres pulling on the nucleus (Fig. 3e). Upon stretching, the pulling forces generated on the nucleus by adhesion progressively diminished due to the nucleus disengaging from the keratin network. By contrast, the steric pressure increased sharply as the fibres tightened against the nucleus, then dropped abruptly after nuclear escape (Fig. 3e). Counterintuitively, this suggests that pulling on the tissue results in nuclear compression, which eventually pushes the nucleus out of its surrounding keratin cage.

### The nucleus is pushed out of its keratin cage

To test this prediction, we first inspected the fixed spontaneous domes (Fig. 1b). As predicted, we found that the star-like bundle structures are localised on the basal side of the cell while the nucleus is positioned on top of this basal structure and is surrounded by only a residual amount of keratin. This stands in stark contrast to non-bundled cells, where the nucleus resides in the medial plane and is surrounded by a prominent keratin cage in all three dimensions (Fig. 1b, Supplementary Fig. 2).

To understand the dynamics of this drastic structural transition, we performed simultaneous live confocal imaging of keratin-18-GFP and the nucleus (labelled with SPY-DNA) at constant dome pressure (Fig. 3f). At the onset of inflation, before bundling, the peaks of fluorescence intensity of keratin and DNA roughly colocalized along the z axis, indicating that the nucleus was fully embedded in its keratin cage. With time, the nucleus progressively escaped from the cage, as revealed by an increasing offset between the keratin and DNA peaks (Fig.3g,h). Eventually, the nucleus fully popped out of the cage, giving rise to the formation of the central knot characteristic of star-like bundled cells (Fig. 3f-h). To study the coupling between the bundling transition and nuclear escape, we plotted the nuclear offset with respect to *θ* and *Ri*. We found that the offset is strongly correlated with both metrics across individual cells, indicating a robust link between keratin bundling and nuclear uncaging (Fig. 3i,j). Together, our computational model and experiments paint a physical picture whereby a tension increase in keratin fibres generates a compressive stress on the nucleus, which is ultimately squeezed out of its cage.

### Supracellular clusters emerge through nucleation and growth

We finally investigated the supracellular dynamics of the bundling transition. Fixed images of spontaneous domes show that bundled cells arrange in multicellular clusters (Fig. 1a,c). These clusters could emerge through different spatiotemporal scenarios. A first possibility is that bundled cells initially form in separate locations of the dome and subsequently move to coalesce into a single cluster. Alternatively, all cells in a cluster could bundle simultaneously, forming a supracellular bundled network in a single step. A third scenario is that bundling nucleates in a single cell and progressively propagates by recruiting neighbouring cells into the cluster.

To distinguish between these different scenarios, we analysed timelapse sequences of keratin18- GFP domes subjected to a step pressure of 200 Pa, ensuring that none of the domes had experienced prior inflation. A representative sequence is shown in Fig. 4a, where the cell outline is colour-coded according to the average angle *θ* of each cell at the corresponding time point (Supplementary Video 4). This sequence shows that, following an initial period without detectable bundling, one cell undergoes a bundling transition while some of its neighbours exhibit a progressive increase in *θ*. With time, these neighbouring cells also cross the bundling threshold, leading to the growth of a cluster. This observation suggests that bundling occurs sequentially rather than simultaneously and proceeds through cluster nucleation and growth rather than coalescence.

**Fig 4:**
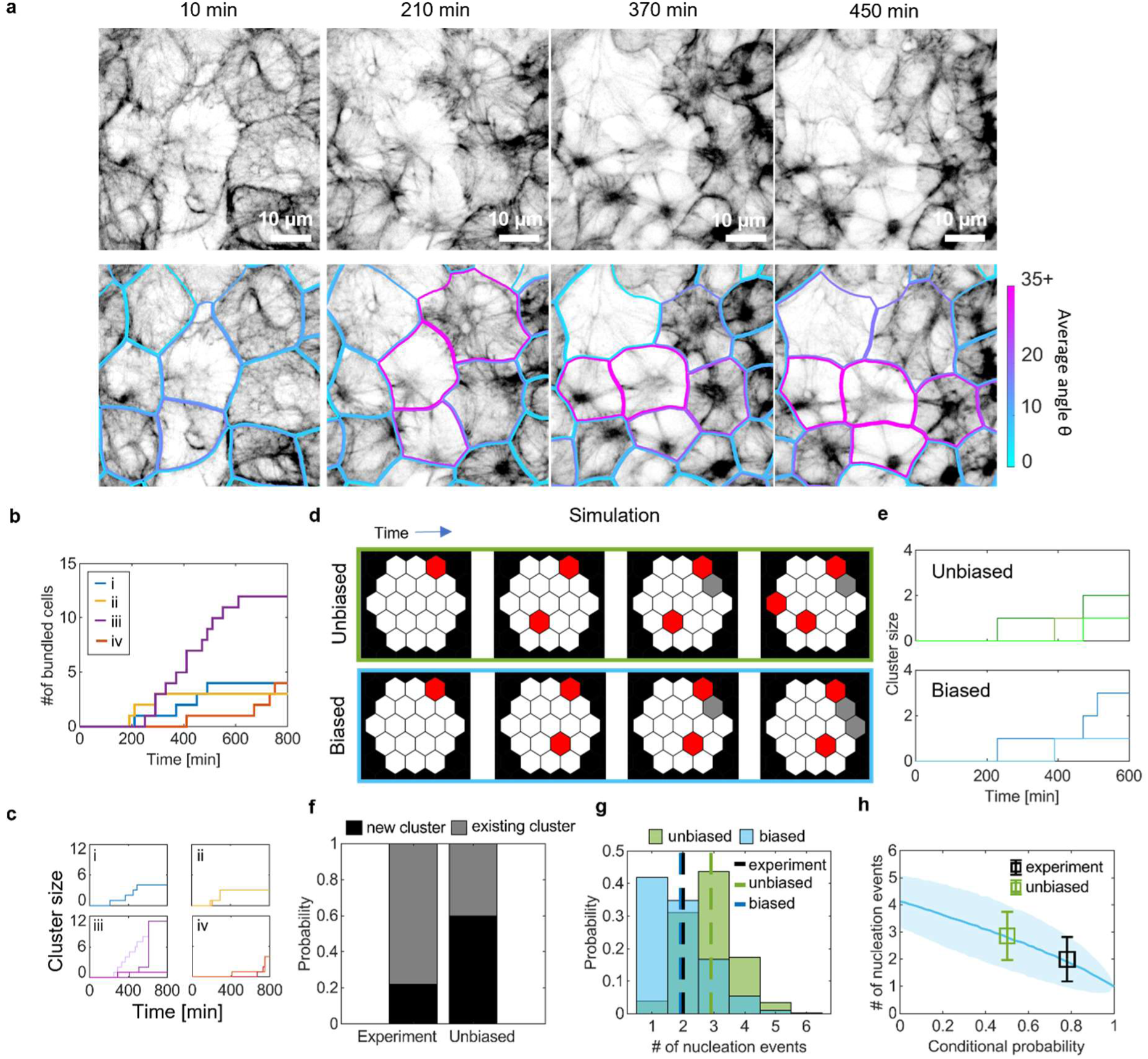
B**u**ndling **is a collective process. a,** Cropped images of MDCK keratin-18-GFP cells in a directed dome. Outlines indicate segmented cell borders color-coded according to the average angle *θ*. Frames are chosen to display individual cells crossing the bundling threshold sequentially. **b**, Number of bundled cells over time in 4 domes. **c**, Cluster size over time in 4 domes. If a dome has multiple clusters, they are shown with different intensities until they merge. **d**, Examples of sequential bundling events in unbiased and biased simulations. Black lines represent cell borders. Cells starting a new cluster count as a nucleation event and are marked in red. Cells bundling adjacent to an existing cluster are marked in grey. **e,** Cluster size over time in the simulations shown in (**d**). **f**, Probability of bundling cells to form a new cluster (black) or join an existing cluster (grey) 100 min after the first bundled cell in a dome. Data are shown for experiments and for the unbiased simulation. **g**, Probability distribution of the number of nucleation events per dome at the end of the simulation. Dashed lines indicate mean values. (n = 4000 simulations each). **h**, Mean number of nucleation events plotted against the conditional probability to join an existing cluster in biased simulations. Shaded area represents standard deviation. Data points show the experimental probability and the unbiased simulation probability, represented as mean ± s.d.

To substantiate this observation quantitatively, we computed the number of bundled cells per dome over time. This analysis, performed over four different domes, confirms that bundling is a sequential process that starts approximately after 200 min of inflation and proceeds at an average rate of 1.00 ± 0.65 bundled cells per hour (mean ± s.d.) (Fig. 4a,b). In addition, our analysis shows that bundling occurs through the nucleation of clusters (2.0 ± 0.8, mean ± s.d. nucleation events per dome) that grow over time and eventually merge into larger ones (Fig. 4c, Supplementary Fig. 8a).

These experiments suggest that a cell is more likely to bundle if at least one of its neighbours has already bundled. Hence, the growth of a cluster is favoured over the creation of a new one. To ascertain this, we performed a Monte-Carlo-simulation of dome dynamics under different probabilistic scenarios for a cell to bundle. In our simulations, domes are modelled as 2D hexagonal grids of 19 cells, matching the experimental dome size (Figs. 4d). Initially, no cells are bundled. The simulation considers sequential bundling events, in which a cell is randomly selected to switch to the bundled state, and once bundled, it remains in that state. In a first unbiased scenario, all non-bundled cells have equal probability of being selected for switching to a bundled state. The times selected for bundling were taken from the experiments to allow for a direct comparison between simulations and observations (Fig. 4e). We found that, within the first 100 minutes, the probability of forming a new cluster was 60%, compared to just 22% in the experiments (Fig. 4f). Both probabilities decreased over time as some clusters had already formed, yet the nearly threefold difference between simulation and experiment persisted (Supplementary Fig. 8b). The average number of nucleation events at the end of the simulation was 2.9 ± 0.9 (mean ± s.d., dashed green line in Fig. 4g), far above the 2 ± 0.8 (mean ± s.d. dashed black line in Fig. 4g)) observed experimentally. Thus, an unbiased scenario where each cell in the dome has an equal probability to bundle does not account for our experimental results.

This discrepancy led us to consider a second biased scenario, in which the probability of bundling is higher if a neighbouring cell had already bundled. As expected, increasing this probability decreased the number of nucleation events in the domes (Fig. 4h). Remarkably, if each cell was given a 78% chance to be selected if it was adjacent to an existing cluster and a 22% chance if it was not, as measured experimentally (black bars in Fig. 4f), the average number of nucleation events was 1.9 ± 0.9 (mean ± s.d., dashed blue line in Fig. 4g), matching the experimental results (dashed black line in Fig. 4g). Together, these simulations support the conclusion that the spatial distribution of bundling events is not completely random but instead arises from a slow nucleation and growth mechanism.

## Discussion

In this work we studied the dynamics of the keratin cytoskeleton during the application of large and sustained stretch. We identified a structural transition that spans multiple length scales. At the subcellular scale, keratin filaments assemble into thick bundles that radiate from a central tangle and bisect cell-cell junctions. At the supracellular scale, cells undergoing this star- bundling transition nucleate clusters that grow in time to form domains. Interestingly, this transition reflects a geometric transformation between two topological representations of the tissue: the tessellation—defined by the tiling of the plane by polygonal cells, where intermediate filaments initially accumulate at cell–cell junctions—and a dual triangulation formed by supracellular bundles connecting central knots. According to this notion of topological duality, polygons in the tessellation (cells) become vertices in the triangulation (knots), whereas edges in the tessellation (cell-cell junctions) become crossing edges in the triangulation (keratin bundles). The bundling transition is dynamically regulated by the keratin-actin interaction and requires the escape of the nucleus from its keratin cage.

The keratin cytoskeleton is now recognized to play a protective role against large deformations. Such role has traditionally been attributed to the strain-stiffening properties of keratin filament networks, which have been linked to structural rearrangements such as keratin bundling ^17,19^. However, our findings reveal that the timescales of strain-stiffening and the bundle transition are fundamentally distinct. While strain-stiffening has been shown to occur over time scales of seconds to minutes^17^, the bundling transition and its supracellular propagation span hours to days. The keratin bundling transition should therefore not be interpreted as the “safety belt” mechanism that provides strain-stiffening to the tissue under large and fast deformations^44^. Rather, the star-bundling transition can be viewed as the terminal stage in a progressive time-dependent reorganization of the keratin network, which tends to relax stresses until topologically possible by the formation of the central knot, shifting the onset of the protective strain-stiffening regime to larger strains.

A remarkable feature of the structural transition uncovered in this study is the complete escape of the nucleus from its keratin cage. This perinuclear cage is generally thought to play a protective role by shielding the nucleus from large mechanical deformations, in line with observations in single breast epithelial cells where a keratin perinuclear network renders the nucleus insensitive to the mechanics of its environment ^10^. Our computational results suggest that, at moderate stretch, the intermediate filament network indeed wraps the nucleus more tightly, shielding it from the applied tension. However, upon large and sustained stretching, both our experiments and simulations show that the nucleus leaves the keratin cage. Whether this uncaging process compromises or protects the nucleus is a major question raised by this study. On the one hand, an unshielded nucleus is more vulnerable to mechanical forces. On the other hand, and perhaps counter-intuitively, decoupling the nucleus from the highly stressed cytoskeleton may provide an alternative form of protection by isolating it from force transmission pathways. This hypothesis could be addressed experimentally in the future, for example, by analyzing nuclear damage under conditions of prolonged mechanical strain.

Our joint experimental and computational analysis identifies the main determinants that govern the onset and dynamics of the star-bundling transition. A first major factor is the interaction between keratin and actin, which contributes to slow down the transition. Consistent with this result, previous work showed that a loss of function of the actin-keratin crosslinker plectin is sufficient to bundle keratin filaments, even in the absence of mechanical stretch ^38^. In addition, large deformations are known to induce thinning of the actin cortex, potentially reducing the number of actin-keratin interactions and thereby favoring the bundling transition ^19^. A second key determinant is the escape of the nucleus from its perinuclear keratin cage. For the star-bundling transition to complete, the keratin network must overcome a mechanical obstacle that arises from the specific and steric interactions with the nucleus. A third potential determinant of the star-bundling transition is the interaction with the extracellular matrix (ECM), either directly through hemidesmosomal junctions or indirectly via focal adhesions. While this interaction cannot be tested in our experimental system, where stretched cells are largely devoid of ECM, previous work from our groups has shown that the adhesion of keratin to the laminin through hemidesmosomes affects cytoskeleton mechanics and shields the nucleus from actomyosin-mediated mechanical deformation, potentially impairing the star-bundling transition ^10^. Interestingly, a recent study in the intestinal epithelium showed that, in conditions of low adhesion, actin can adopt a star-shape morphology similar to the one observed here for keratin ^45^. These results suggest that the star-shape organization of cytoskeleton filaments might be a general mechanical adaptation, although it is unclear if the underlying mechanisms and dynamics of actin star formation are analogous to those observed here for keratin.

By uncovering the mechanisms underlying the star-like bundling transition, our study identifies the conditions under which this transition may be relevant in physiology and disease. In early development, tissues are subjected to slow and sustained deformations while often expressing relatively low levels of ECM—two factors that favor the bundling transition. It will therefore be interesting to investigate whether keratin undergoes partial or complete bundling transitions in developmental systems such as inflating blastocysts or gastrulating vertebrate embryos ^46,47^. In adulthood, the bundling transition might be relevant in glandular organs experiencing large and sustained stretching, such as the bladder or the lactating mammary gland. In the context of disease, keratin bundling transitions may play a role in conditions where keratin filament interactions are impaired, such as in epidermolysis bullosa simplex (EBS)^48^. Beyond potential physiological implications, the physics underlying the star-bundling transition could inform the design of bioinspired polymeric materials. By harnessing the mechanisms identified in this study, such materials could be engineered to display transitions between states with markedly distinct material properties, along with controllable transition times and operating length scales.

## Methods

### Cell culture

MDCK strain II cells were used throughout the study. To visualize intermediate filaments, MDCK cells expressing keratin-18-GFP were used^29^. To visualize the plasma membrane, MDCK keratin- 18-GFP co-expressing CAAX-iRFP were used. To test the effects of plectin MDCK keratin-18-GFP cells co-expressing P1f-8-BFP were used. All MDCK cell lines were cultured in Dulbecco’s Modified Eagle Medium (DMEM, Gibco) with high glucose and L-glutamine supplemented with with 10% v/v fetal bovine serum (FBS, Gibco), 100 µg/ml penicillin and 100 µg/ml streptomycin. Cells were maintained at 37°C in a humidified atmosphere with 5% CO2. Live imaging of nuclei was performed by incubating cells (12 h, 1/1000 dilution) with SPY650-DNA (Spirochrome). MDCK keratin-18-GFP cells were obtained from the laboratory of G. Charras. MDCK CAAX-iRFP cells were obtained via transposon insertion of iRFP-CAAX. MDCK P1f-8-BFP were obtained via transposon insertion of P1f-8-BFP. The P1f-8-BFP cassette was generated by replacing the EGFP of the plasmid pGR240^38,49^ by the EBFP2 of the Addgene plasmid #55242, via PCR and Gibson assembly. All cell lines were tested negative for mycoplasma contamination.

### Fabrication of soft silicone gels

Soft silicone gels were prepared based on previously published protocols ^50,51^. Briefly, silicone elastomer was synthesized by mixing a 1:1 weight ratio of CY52-276A and CY52-276B polydimethylsiloxane (PDMS, Dow Corning Toray). After degassing for 30 min on ice, 120 µl of the gel was spin-coated on glass-bottom dishes (35 mm, no. 0 coverslip thickness, Mattek) for 10 s at 500 rpm and for 40 s at 5000 rpm. The samples were cured at 65°C overnight.

### Coating of soft PDMS gels with fluorescent beads

Previously cured soft PDMS gels were treated with (3-aminopropyl)triethoxysilane (APTES, Sigma-Aldrich) diluted at 5% in absolute ethanol for 3 min, rinsed 3 times with 96% ethanol, and dried in to the oven for 30 min at 60°C. Fluorescent carboxylate-modified beads (FluoSpheres dark red, invitrogen) were filtered (220 nm) and sonicated in sodium tetraborate (3.8 mg/ml, Sigma-Aldrich), boric acid (5 mg/ml, Sigma-Aldrich), and 1-ethyl-3-(3-dimethylaminoprpyl)carbodiimide (EDC, 0.1 mg/ml, Sigma-Aldrich), as described previously ^51,52^. The bead solution was placed for 5 min on the gels and then rinsed 3 times with type-1 water. The beads were passivated by incubation with tris-buffered saline (TBS, Sigma-Aldrich) for 20 min at room temperature, rinsed 3 times with type-1 water and dried in the oven for 15 min at 60°C.

### Micropatterning of soft PDMS gels

Stamps for micropatterning dome footprints were fabricated from SU8-50 masters containing 100 µm diameter cylinders that were raised by conventional photolithography. Uncured PDMS (Sylgard, Dow Corning) was poured on the masters and cured overnight at 65°C. The cured PDMS was peeled of and cut into individual stamps. For the patterning, the PDMS stamps were incubated with 40 µg/ml fibronectin (fibronectin from human plasma, Sigma-Aldrich) and 20 µg/ml Fibrinogen Alexa Fluor 647 Conjugate (Invitrogen) in phosphate-buffered saline (PBS, Sigma-Aldrich) for 1h and then brought into contact with poly vinyl alcohol (PVA, Sigma Aldrich) membranes to transfer the protein. The PVA membranes were gently placed on the soft PDMS gels for 1 h and dissolved in a 0.2% w/v solution of Pluronic F127 (Sigma-Aldrich) overnight to passivate the non-patterned surface at the same time. Afterwards, the gels were washed with PBS.

### Fabrication of microfluidic pressure device

A detailed protocol for fabricating the microfluidic device for applying hydrostatic pressure on cell monolayers can be found here^53^. It consists of five parts. From top to bottom, the first part is a thick PDMS block with four inlets and a channel connected to two inlets on its bottom. The second is a 100 µm thin PDMS layer with a 5x5 square pattern of 80 µm diameter holes and two inlets passing through that layer. The third part is polycarbonate membrane with 400 nm sized holes (Whatman Nuclepore Track-Etch Membrane 0.4 µm). The fourth part is a roughly 120 µm thick PDMS layer with a channel perpendicular to the upper block channel and the fifth part is a glass-bottom dish (35 mm dish #0 glass, Cellvis).

The top block was made via replica molding in a 3D printed mold from vat polymerization with a digital light processing 3D printer (Solus DLP 3D Printer with SolusProto resin). The surface of the mold was silanized to prevent adhesion with unpolymerized PDMS. PDMS was mixed at a 1:9 curing agent to elastomer weight ratio and degassed for 30 min while the mold was preheated to 90°C. The degassed PDMS was then poured into the mold, degassed again for 30 min and cured on a hot plate at 90°C for at least 30 min. After curing, the PDMS was gently removed from the mold, cut into individual blocks, and punched with a 1.5 mm Rapid-Core Sampling Tool (Electron Microscopy Sciences) to generate four inlets. The second PDMS part was fabricated from SU8-50 masters containing 80 µm diameter columns. For one wafer, 10 ml elastomer was mixed with 1 ml curing agent, degassed for 30 min, and distributed on the wafer in way that covers all features. The wafer was then spun at 600 rpm for 1 min in a spin coater (WS-650MZ 23NPP/LITE, Laurell Tech), an air gun was used to gently remove uncured PDMS from the top of all features. The wafer was then kept on a levelled hot plate over night at room temperature before curing at 90°C for 30 min the next morning. The top block and this part were connected by treating both pieces for 1 min with plasma in an ozone plasma cleaner (PCD-002-CE, Harrick Plasma). Afterwards, they were brought into direct contact with each other and bonded for 2 h at 80°C.

The bottom layer with one channel was made by mixing PDMS in a 2:8 curing agent to elastomer ratio and pouring 3.5 ml on a plastic dish (tissue culture dish 150, TPP), distributing and removing air bubbles with an air gun, keeping the plastic dish on a levelled surface overnight at room temperature, and curing for at least 1 h at 80°C. After curing, the thin PDMS sheets were placed on a Silhouette cutting mat and cut into individual pieces on a Silhouette cutting machine (Silhouette Cameo 4, Silhouette America). The resulting pieces were connected to the glass bottom dish via plasma bonding for 1 min and resting in an oven for 2 h at 80°C.

The polycarbonate membrane was cut into small pieces big enough to cover the intersection of two perpendicular channels and activated in a plasma cleaner for 1 min at 600 mTorr. They were then covered by a preheated 80°C warm mixture of 5% APTES in MilliQ water for 20 min before drying on cleanroom tissue. These coated membranes were bonded to the top block by plasma cleaning the block for 20 s at 600 mTorr and bringing the previously plasma treated side of the membrane in contact with the block. The remaining two pieces, the top block with the thin PDMS layer and the membrane plus the glass bottom dish with the bottom channel PDMS layer, are finally connected by plasma cleaning both parts for 30 s and gently pressing them together without breaking the glass of the glass bottom dish. To increase stability, the base of the resulting device was surrounded by uncured PDMS and placed in the oven for 2 h at 80°C.

### Patterning of microfluidic pressure device

Surface protein patterns were applied inside the microfluidic devices via light-induced molecular adsorption of proteins with PRIMO (Alveole). The devices were first filled with 96% ethanol to remove bubbles and then washed with MilliQ water. Afterwards, they were incubated with Poly-L-lysine solution (PLL, Merck) for at least 1 h at room temperature. Next, they were washed with HEPES buffer (pH 8.4) and incubated with a freshly prepared solution of mPEG-Succinimidyl Valederate (SVA-PEG, Laysan Bio) in the same HEPES buffer at 50 mg/ml overnight at 4°C. Directly before using PRIMO, the devices were washed with MilliQ water and incubated with the photo initiator (4-Benzoylbenzyl)trimethylammonium chloride (PLPP, Angene). The actual patterned was applied with PRIMO by illuminating the sample with a UV-laser using the highest intensity on the area surrounding the dome foodprints and using 0% or 10 % of the intensity on the foodprints. The UV treated devices were then washed with PBS and incubated with an ice-cold solution of fibronectin (100 µg/ml) and fibrinogen Alexa Fluor 647 (30 µg/ml) in PBS for 5 min. After washing with PBS, the devices could be used immediately for cell seeding or stored overnight a 4°C.

### Cell seeding

Samples were sterilized for 15 min under UV and washed with sterile PBS directly before seeding. For spontaneous domes on soft PDMS, 80 µl of cells in medium at a concentration of 2 000 000 cells/ml were placed on the gel, incubated for 50 min, washed with PBS to remove non-attached cells, and incubated with 2 ml of Eagle’s minimal essential medium (MEM, Gibco) supplemented with 10% v/v fetal bovine serum (FBS, Gibco), 100 µg/ml penicillin and 100 µg/ml streptomycin.

For directed domes in microfluidic devices, 35 µl of cells in medium at a concentration of 30 000 000 cells/ml were added to the bottom channel and incubated for 50 min. Afterwards, the samples were washed with medium to remove unattached cells. If the seeding did not result in a confluent cell layer with the dome footprints being mostly unoccupied, the seeding could be repeated once with the same parameters. The samples were then incubated for 24 h before the experiment.

### Directed dome experiments

One day before the experiment, a reservoir of CO2 independent medium (ThermoFisher) supplemented with 10% v/v fetal bovine serum (FBS, Gibco), 100 µg/ml penicillin, 100 µg/ml streptomycin, and 1.3% v/v HEPES solution(Sigma-Aldrich) was incubated overnight at 37°C and degassed for at least 30 min the next morning to remove all dissolved gas from the medium and prevent bubble formation in the tubes during the experiment. The cell medium in the microfluidic device was replaced by the same CO2 independent medium. After mounting the microfluidic chip to the stage of the temperature-controlled microscope, the reservoir of CO2 independent medium was connected with a silicone tube to the inlet of the pressure channel. A second silicon tube with a valve was then connected to the outlet of the pressure channel. A weak vacuum was connected to the outlet valve to remove all bubbles in the tubes and the outlet valve was closed.

Hydrostatic pressure was induced by increasing the hight of the reservoir relative to the height of the microfluidic device with the help of a vertically mounted motorized stage. The stage height for zero pressure was determined by increasing height until domes formed and gradually lowering the stage height until domes disappeared, thus marking 0 Pa. The hydrostatic pressure during the experiments was determined with the conversion of 1 𝑚𝑚 = 10 𝑃𝑎 based on Pascals law.

For LatrunculinA experiments, 1µM of LatrunculinA (Sigma) dissolved in 200 µl of CO2 independent medium was added gently with a pipette directly into the cell channel.

### Immunostainings

Spontaneous domes were fixed with 4 % paraformaldehyde in PBS for 10 min at room temperature and permeabilized with 0.1 % Triton X100 (Sigma-Aldrich) in PBS for 10 min at room temperature. Afterwards a blocking solution of 1 % bovine serum albumin (BSA, Sigma-Aldrich) in PBS was used for 1 h at room temperature. F-actin was labelled by adding tetramethylrhodamine phalloidin (TRITC phalloidin, Sigma-Aldrich) at 1:1000 dilution in PBS for 30 min at room temperature. Nuclei were labelled by adding Hoechst (Hoechst 33342, invitrogen) at 1:2500 dilution in PBS for 10 min.

Imaging was performed on a Zeiss LSM880 confocal microscope running the software ZEISS2.3SP1FP3(black, version 14.0.24.201) using the FastAiryScan mode and a Plan-Apochromate 63x 1.4-numerical aperture oil immersion objective. Dome footprint positions were obtained with a 631 nm laser scan to visualize the fibrinogen Alexa Fluor 647 conjugate. Three channels were acquired: 405 nm to excite Hoechst, 488 nm to excite keratin-18-GFP, and 561 nm to excite tetramethylrhodamine phalloidin. Z-stacks of all three channels were obtained sequentially with a step size of 144 nm.

### Time-lapse microscopy

Experiments were carried out on a Zeiss LSM880 confocal microscope running the software ZEISS2.3SP1FP3(black, version 14.0.24.201) using the FastAiryScan mode. For the live directed domes, imaging was performed with a LD-LCI Plan-Apochromat 40x 1.2-numerical aperture water immersion objective or a Plan-Apochromate 63x 1.4-numerical aperture oil immersion objective. Dome footprint positions were obtained with a 561 nm laser scan to visualize the fibrinogen Alexa Fluor 546 conjugate. Depending on the experiment, the following channels were acquired: 488 nm to excite keratin-18-GFP, 631 nm to excite SPY-650 DNA, 631 nm to excite iRFP-CAAX, and 405 nm to excite P1f-8-BFP. To prevent photodamage, the 405 nm channel was only used in the first and the last time point to provide a reference. Z-stacks were acquired with a 500 nm step size and different channels were acquired sequentially.

### Segmentation and tracking

Cell segmentation was performed with Cellpose ^54^. In case of the fixed spontaneous domes, cells were segmented based on the maximum projection of the actin staining and the cyto1 model. In case of live directed domes, cells were segmented based on the maximum projection of the membrane marked and a cyto2 based model with human-in-the-loop retraining ^55^. All resulting label maps were checked and, if necessary, corrected by hand. In the absence of a membrane or actin signal, cells were segmented by hand based on the keratin signal.

Tracking was performed with the ImageJ plugin trackmate ^56^ based on the label maps. All tracks were checked and, if necessary, corrected by hand.

### Image analysis

Measurement of *Ri* and *θ* was performed on images of the keratin signal of individual cells after cropping based on the segmentation results (see Supplementary Fig. 3). All analysis was performed in Matlab 2020b, unless specified otherwise. The cell centre was determined as the centre of mass of fluorescent intensity. From this centre, the image was transformed into polar coordinates generating a rectangular matrix where each row has the same distance to the pole and each column has the same polar angle. To generate a perfect rectangle and avoid analysis artifacts at the cell edge, all columns were interpolated so that the cell edge reaches the full length of the column. This essentially transforms the cell shape into a circle.

From the obtained rectangular matrix, we measured the mean intensity of each row, yielding an intensity profile as a function of radial distance from the cell centre. We then rescaled the radius by its maximum and calculated *Ri* as the median of the resulting distribution. This resulted in a dimensionless value ranging from 0 to 1, where values near 1 indicate that keratin structures are predominantly localized at the cell periphery, and values near 0 indicate central accumulation.

To obtain *θ*, the same rectangular matrix from the previous step was cropped to contain only the outer half. The resulting image was imported into ImageJ and smoothed with a gaussian blur (σ = 4 px). We then measured the orientation of all keratin structures with the ImageJ plugin OrientationJ ^57^ by calculating the Vector Field (tensor = 8, gradient = 0, vectorgrid = 1) resulting in a local angle and coherency for every pixel ^58^. *θ* was then calculated as the median of the absolute values of all local angles (since they are symmetric around 0°) weighted by their coherency. This led to a single value between 0°, indicating that all keratin structures are parallel to the cell edge, and 90°, indicating that all keratin structures are perpendicular to the cell edge.

Both *Ri* and *θ* were used in the live data to differentiate between bundled and non-bundled cells. For all cells, the maximum average angle *θmax* and the corresponding intensity radius *Rcorr* at the same time point were measured. Cells were marked as being bundled if 𝜃_max_ > 35*°* and 𝑅_corr_ < 0.45, with slight adjustments of *θmax* to avoid false positive or false negative detections. The same threshold values were then used to determine the bundling time *TB. TB* was determined as the first time point, where both *Ri* and *θ* crossed their respective thresholds. In fixed data (Fig. 1), cells were classified as bundled, partly bundled, and non-bundled via visual inspection.

The same analysis was performed for the 2.5D simulations by using the simulated images of the top view as input.

To obtain the cell area in domes, the dome shape was measured with the ImageJ plugin Local Z Projector ^59^, resulting in a height map. The label maps from the segmentation were converted into masks in ImageJ and the actual cell area was calculated from the hight map and the cell masks with DeProj ^59^ in Matlab.

The offset of the nucleus in reference to keratin was obtained from cropped 3D stacks of individual cells containing both the keratin-18-GFP and the SPY650-DNA signal. The stacks were rotated in ImageJ, if necessary, so that the basal plane of the cell was parallel to the image plane. The central plane of the nucleus was then fitted with an ellipse and the keratin signal was cropped in an area that had twice the short and long axis length of the fitted ellipse with the same centre to avoid artifacts from cells with a curved basal plane. The sum of the signal of both channels of the rotated and cropped stacks was then measured as a function of the height in Matlab and the offset of the nucleus in reference to keratin intermediate filaments was calculated as the distance between the respective peaks. The final nuclear offset was then determined over time as a moving mean with window size 5.

### Calculation of tissue strain

Dome strain, defined as 𝜀_D_ = 𝐴_Dome_⁄𝐴_Base_ − 1, was obtained by measuring the radius of curvature 𝑅_C_ and dome height 𝐻 from the side view of the centre of the dome. The dome strain was then calculated as 𝜀_D_ = 2𝑅_C_⁄(2𝑅_C_ − 𝐻) − 1. The constitutive strain 𝜀_C_ was obtained by measuring the mean cell area inside the dome footprint 𝑎_!N_ and the mean cell area outside the dome footprint 𝑎_OUT_ before inflation. The constitutive strain was then calculated via 𝜀_C_ = 𝑎_!N_⁄𝑎_OUT_ − 1 . Finally, the tissue strain 𝜀_T_ was calculated as the combination of both strains via 𝜀_T_ = 𝜀_D_ + 𝜀_C_ + 𝜀_D_𝜀_C_ . This calculation accounts for the fact that, before inflation, cells inside the footprint had a larger area than those outside of it, as a consequence of sample preparation.

### Monte-Carlo-simulations

Monte-Carlo-simulations were performed in Matlab R2020b. We first generated a hexagonal grid with 19 cells and a next neighbour list based on shared interfaces. The bundling times of 4 different domes as presented in Fig. 4b were used as input parameters. Each dome was simulated 1000 times, resulting in a total of 4000 simulations. At the start of the simulation, all cells were in a non-bundled state. Using the bundling times in sequence, non-bundled cells were selected to switch into a bundled state until all bundling times of the dome were used. In the unbiased scenario, this selection was performed randomly with an equal chance for every non-bundled cell to be selected. Consequently, bundled cells could not be selected again and stayed in the bundled state. During the simulation, the time and position of each bundled cell was logged. After the simulation, the number of nucleation events of each dome was determined by counting how many cells bundled without having previously bundled neighbours with the help of the next neighbour list.

In the biased scenario, cells were divided into two populations: cells being neighbours of already bundled cells and free cells without a connection to already bundled cells. These populations were updated after every bundling event with the help of the next neighbour list. A conditional probability to select cells from the neighbour population was used as an input parameter with values between 0 and 1. During the simulation, a random number between 0 and 1 was generated for every bundling event. If this number was below the conditional probability, a random cell from the neighbour pool was selected. If the number was above the conditional probability, a random cell from the free pool was selected. If one of the pools had no elements left, a random cell from the other pool was selected. The number of nucleation events was determined in the same way as in the unbiased scenario.

## Statistics

Comparison between each group of unpaired samples were computed using the unpaired two-sided Wilcoxon rank sum test. Correlation was tested by computing the Pearson correlation coefficient R. Statistical analysis was performed with Matlab R2020b.

## Acknowledgements

We thank all the members of our groups for their discussions and support. We thank Mónica Purciolas, Juan Francisco Abenza and Guillermo Martínez Ara for technical assistance. T.G. is funded by the Deutsche Forschungsgemeinschaft (DFG, German Research Foundation) – 445510097. P.R.-C. acknowledges funding from the Spanish Ministry of Science and Innovation (PID2022-142672NB-I00), the European Research Council (AdG 101097753), the Generalitat de Catalunya (2017-SGR-1602), and the prize ‘ICREA Academia’ for excellence in research. X.T. acknowledges funding from the Generalitat de Catalunya (AGAUR SGR-2017-01602), the CERCA Programme, the Spanish Ministry for Science and Innovation MICCINN/FEDER (PID2021- 128635NB-I00 MCIN/AEI/ 10.13039/501100011033 and “ERDF-EU A way of making Europe”), European Research Council (Adv-883739), Fundació la Marató de TV3 (project 201903-30-31-32), European Commission (H2020-FETPROACT-01-2016-731957), La Caixa Foundation (LCF/PR/HR24/00326) and the Human Frontiers Science Program (HFSPRGP022/2024); IBEC is recipient of a Severo Ochoa Award of Excellence from the MINECO.

## Author contributions

T.G., M.A., X.T. conceived the study. T.G, N.C. and X.T. designed experiments. T.G. performed the experiments and analysed the data. M.P. and M.A. developed the single cell models. T.G. developed the Monte-Carlo simulations. P.R.-C, G.W., G.C., M.A. contributed to material, reagents, technical expertise and discussion. T.G. and X.T. wrote the manuscript. All authors revised the completed manuscript.

## Competing interests

The authors declare no competing financial interests.

## Code availability

Analysis procedures and code implementing the model are available from the corresponding authors on reasonable request.

## Data availability

The data that support the findings of this study are available from the corresponding authors on reasonable request. Extended Data is available for this paper.

## Notes

### Competing Interest Statement

The authors have declared no competing interest.

## References

1. Kreplak, L., Bär, H., Leterrier, J. F., Herrmann, H. & Aebi, U. Exploring the Mechanical Behavior of Single Intermediate Filaments. J. Mol. Biol. 354, 569–577 (2005).

2. Block, J. et al. Nonlinear Loading-Rate-Dependent Force Response of Individual Vimentin Intermediate Filaments to Applied Strain. Phys. Rev. Lett. 118, 048101 (2017).

3. Storm, C., Pastore, J. J., MacKintosh, F. C., Lubensky, T. C. & Janmey, P. A. Nonlinear elasticity in biological gels. Nature 435, 191–194 (2005).

4. Golde, T. et al. The role of stickiness in the rheology of semiflexible polymers. Soft Matter 15, 4865–4872 (2019).

5. Lorenz, C., Forsting, J., Style, R. W., Klumpp, S. & Köster, S. Keratin filament mechanics and energy dissipation are determined by metal-like plasticity. Matter 6, 2019–2033 (2023).

6. Guo, M. et al. The Role of Vimentin Intermediate Filaments in Cortical and Cytoplasmic Mechanics. Biophys. J. 105, 1562–1568 (2013).

7. Mendez, M. G., Restle, D. & Janmey, P. A. Vimentin Enhances Cell Elastic Behavior and Protects against Compressive Stress. Biophys. J. 107, 314–323 (2014).

8. Seltmann, K., Fritsch, A. W., Käs, J. A. & Magin, T. M. Keratins significantly contribute to cell stiffness and impact invasive behavior. Proc. Natl. Acad. Sci. 110, 18507– 18512 (2013).

9. Ramms, L. et al. Keratins as the main component for the mechanical integrity of keratinocytes. Proc. Natl. Acad. Sci. 110, 18513–18518 (2013).

10. Kechagia, Z. et al. The laminin–keratin link shields the nucleus from mechanical deformation and signalling. Nat. Mater. 22, 1409–1420 (2023).

11. Beedle, A. E. M., et al. Fibrillar Adhesion Dynamics Govern the Timescales of Nuclear Mechano-Response via the Vimentin Cytoskeleton. http://biorxiv.org/lookup/doi/10.1101/2023.11.08.566191(2023) doi:10.1101/2023.11.08.566191.

12. Conboy, J. P., Lettinga, M. G., Boukany, P. E., MacKintosh, F. C. & Koenderink, G. H. Actin and vimentin jointly control cell viscoelasticity and compression stiffening. 2025.01.01.630993 Preprint at 10.1101/2025.01.01.630993 (2025).

13. Harris, A. R., Daeden, A. & Charras, G. T. Formation of adherens junctions leads to the emergence of a tissue-level tension in epithelial monolayers. J Cell Sci 127, 2507–2517 (2014).

14. Price, A. J. et al. Mechanical loading of desmosomes depends on the magnitude and orientation of external stress. Nat. Commun. 9, 5284 (2018).

15. Kröger, C. et al. Keratins control intercellular adhesion involving PKC-α–mediated desmoplakin phosphorylation. J Cell Biol 201, 681–692 (2013).

16. Hatzfeld, M., Keil, R. & Magin, T. M. Desmosomes and Intermediate Filaments: Their Consequences for Tissue Mechanics. Cold Spring Harb. Perspect. Biol. 9, a029157 (2017).

17. Duque, J. et al. Rupture strength of living cell monolayers. Nat. Mater. 23, 1563–1574 (2024).

18. Wang, N., Butler, J. P. & Ingber, D. E. Mechanotransduction Across the Cell Surface and Through the Cytoskeleton. Science 260, 1124–1127 (1993).

19. Latorre, E. et al. Active superelasticity in three-dimensional epithelia of controlled shape. Nature 563, 203 (2018).

20. Huber, F., Boire, A., López, M. P. & Koenderink, G. H. Cytoskeletal crosstalk: when three different personalities team up. Curr. Opin. Cell Biol. 32, 39–47 (2015).

21. Wiche, G. Plectin-Mediated Intermediate Filament Functions: Why Isoforms Matter. Cells 10, 2154 (2021).

22. Wiche, G., Osmanagic-Myers, S. & Castañón, M.J. Networking and anchoring through plectin: a key to IF functionality and mechanotransduction. Curr. Opin. Cell Biol. 32, 21–29 (2015).

23. Russell, D., Andrews, P. D., James, J. & Lane, E. B. Mechanical stress induces profound remodelling of keratin filaments and cell junctions in *epidermolysis bullosa simplex* keratinocytes. J. Cell Sci. 117, 5233–5243 (2004).

24. Ridge, K. M. et al. Keratin 8 Phosphorylation by Protein Kinase C δ Regulates Shear Stress-mediated Disassembly of Keratin Intermediate Filaments in Alveolar Epithelial Cells*. J. Biol. Chem. 280, 30400–30405 (2005).

25. Kreplak, L. & Fudge, D. Biomechanical properties of intermediate filaments: from tissues to single filaments and back. BioEssays 29, 26–35 (2007).

26. Fois, G. et al. Effects of keratin phosphorylation on the mechanical properties of keratin filaments in living cells. FASEB J. 27, 1322–1329 (2013).

27. Lutz, A. et al. Acute effects of cell stretch on keratin filaments in A549 lung cells. FASEB J. 34, 11227–11242 (2020).

28. Meyer, R. et al. The keratin cortex stabilizes cells at high strains. Preprint at 10.1101/2025.02.24.639846 (2025).

29. Harris, A. R. et al. Characterizing the mechanics of cultured cell monolayers. Proc. Natl. Acad. Sci. 109, 16449–16454 (2012).

30. Nahaboo, W. et al. Keratin filaments mediate the expansion of extra-embryonic membranes in the post-gastrulation mouse embryo. EMBO J. 41, e108747 (2022).

31. Leighton, J., Brada, Z., Estes, L. W. & Justh, G. Secretory Activity and Oncogenicity of a Cell Line (MDCK) Derived from Canine Kidney. Science 163, 472–473 (1969).

32. Tanner, C., Frambach, D. A. & Misfeldt, D. S. Transepithelial transport in cell culture. A theoretical and experimental analysis of the biophysical properties of domes. Biophys. J. 43, 183–190 (1983).

33. Quinlan, R. A. et al. A rim-and-spoke hypothesis to explain the biomechanical roles for cytoplasmic intermediate filament networks. J. Cell Sci. 130, 3437–3445 (2017).

34. Windoffer, R. et al. Quantitative mapping of keratin networks in 3D. eLife 11, e75894 (2022).

35. Pensalfini, M., Golde, T., Trepat, X. & Arroyo, M. Nonaffine Mechanics of Entangled Networks Inspired by Intermediate Filaments. Phys. Rev. Lett. 131, 058101 (2023).

36. Strale, P.-O. et al. Multiprotein Printing by Light-Induced Molecular Adsorption. Adv. Mater. 28, 2024–2029 (2016).

37. Broussard, J. A. et al. Scaling up single-cell mechanics to multicellular tissues – the role of the intermediate filament–desmosome network. J. Cell Sci. 133, (2020).

38. Prechova, M. et al. Plectin-mediated cytoskeletal crosstalk controls cell tension and cohesion in epithelial sheets. J. Cell Biol. 221, e202105146 (2022).

39. Moch, M. & Leube, R. E. Hemidesmosome-Related Keratin Filament Bundling and Nucleation. Int. J. Mol. Sci. 22, 2130 (2021).

40. Andrä, K., Nikolic, B., Stöcher, M., Drenckhahn, D. & Wiche, G. Not just scaffolding: plectin regulates actin dynamics in cultured cells. Genes Dev. 12, 3442–3451 (1998).

41. Nikolic, B., Mac Nulty, E., Mir, B. & Wiche, G. Basic amino acid residue cluster within nuclear targeting sequence motif is essential for cytoplasmic plectin-vimentin network junctions. J. Cell Biol. 134, 1455–1467 (1996).

42. Nedelec, F. & Foethke, D. Collective Langevin dynamics of flexible cytoskeletal fibers. New J. Phys. 9, 427 (2007).

43. Wilhelmsen, K. et al. Nesprin-3, a novel outer nuclear membrane protein, associates with the cytoskeletal linker protein plectin. J. Cell Biol. 171, 799–810 (2005).

44. Qin, Z., Kreplak, L. & Buehler, M. J. Hierarchical Structure Controls Nanomechanical Properties of Vimentin Intermediate Filaments. PLOS ONE 4, e7294 (2009).

45. Barai, A. et al. A multicellular actin star network underpins epithelial organization and connectivity. 2024.07.26.605277 Preprint at 10.1101/2024.07.26.605277 (2024).

46. Dumortier, J. G. et al. Hydraulic fracturing and active coarsening position the lumen of the mouse blastocyst. Science 365, 465–468 (2019).

47. Chan, C. J. et al. Hydraulic control of mammalian embryo size and cell fate. Nature 571, 112–116 (2019).

48. McMillan, J. R., McGrath, J. A., Tidman, M. J. & Eady, R. A. J. Hemidesmosomes Show Abnormal Association with the Keratin Filament Network in Junctional Forms of Epidermolysis Bullosa. J. Invest. Dermatol. 110, 132–137 (1998).

49. Rezniczek, G. A., Abrahamsberg, C., Fuchs, P., Spazierer, D. & Wiche, G. Plectin 5′- transcript diversity: short alternative sequences determine stability of gene products, initiation of translation and subcellular localization of isoforms. Hum. Mol. Genet. 12, 3181–3194 (2003).

50. Vedula, S. R. K. et al. Emerging modes of collective cell migration induced by geometrical constraints. Proc. Natl. Acad. Sci. 109, 12974–12979 (2012).

51. Marín-Llauradó, A. et al. Mapping mechanical stress in curved epithelia of designed size and shape. Nat. Commun. 14, 4014 (2023).

52. W. Style, R., et al. Traction force microscopy in physics and biology. Soft Matter 10, 4047–4055 (2014).

53. Chahare, N. et al. Multiscale wrinkling dynamics in epithelial shells. 2025.06.30.662426 Preprint at 10.1101/2025.06.30.662426 (2025).

54. Stringer, C., Wang, T., Michaelos, M. & Pachitariu, M. Cellpose: a generalist algorithm for cellular segmentation. Nat. Methods 18, 100–106 (2021).

55. Pachitariu, M. & Stringer, C. Cellpose 2.0: how to train your own model. Nat. Methods 19, 1634–1641 (2022).

56. Ershov, D. et al. TrackMate 7: integrating state-of-the-art segmentation algorithms into tracking pipelines. Nat. Methods 19, 829–832 (2022).

57. Püspöki, Z., Storath, M., Sage, D. & Unser, M. Transforms and Operators for Directional Bioimage Analysis: A Survey. in *Focus on Bio-Image Informatics* (eds. De Vos, W. H., Munck, S. & Timmermans, J.-P.) 69–93 (Springer International Publishing, Cham, 2016). doi:10.1007/978-3-319-28549-8_3.

58. Rezakhaniha, R. et al. Experimental investigation of collagen waviness and orientation in the arterial adventitia using confocal laser scanning microscopy. Biomech. Model. Mechanobiol. 11, 461–473 (2012).

59. Herbert, S. et al. LocalZProjector and DeProj: a toolbox for local 2D projection and accurate morphometrics of large 3D microscopy images. BMC Biol. 19, 136 (2021).

