## Supplementary Information for "A keratin bundling transition uncages the nucleus in stretched epithelia"

This file contains:

Supplementary figures 1-8

Supplementary video captions

Supplementary note

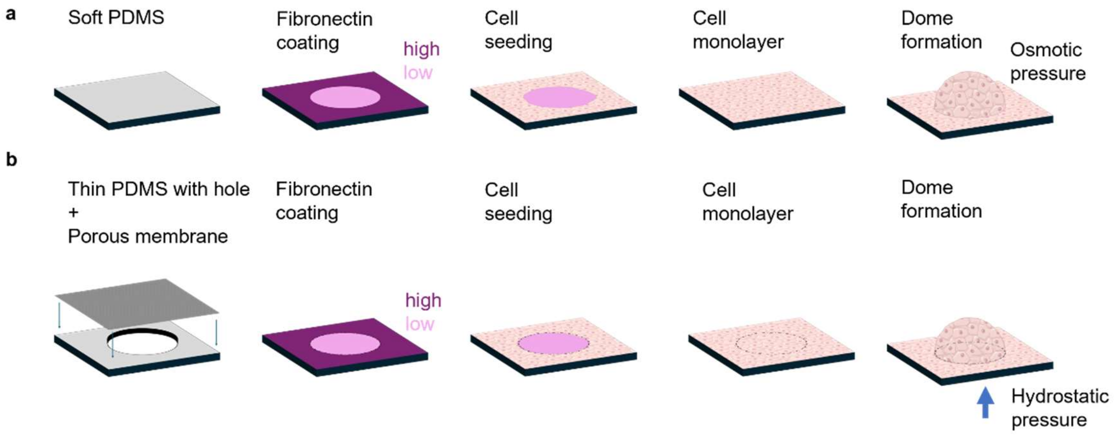


**Supplementary Figure 1: Experimental design of spontaneous and directed domes. a,** Illustration of the different steps in preparing micropatterned spontaneous domes. **b,** Illustration of different steps in preparing micropatterned directed domes. See methods section for details. Monolayer illustration adapted from NIAID NIH BIOART Source (bioart.niaid.nih.gov/bioart/240).

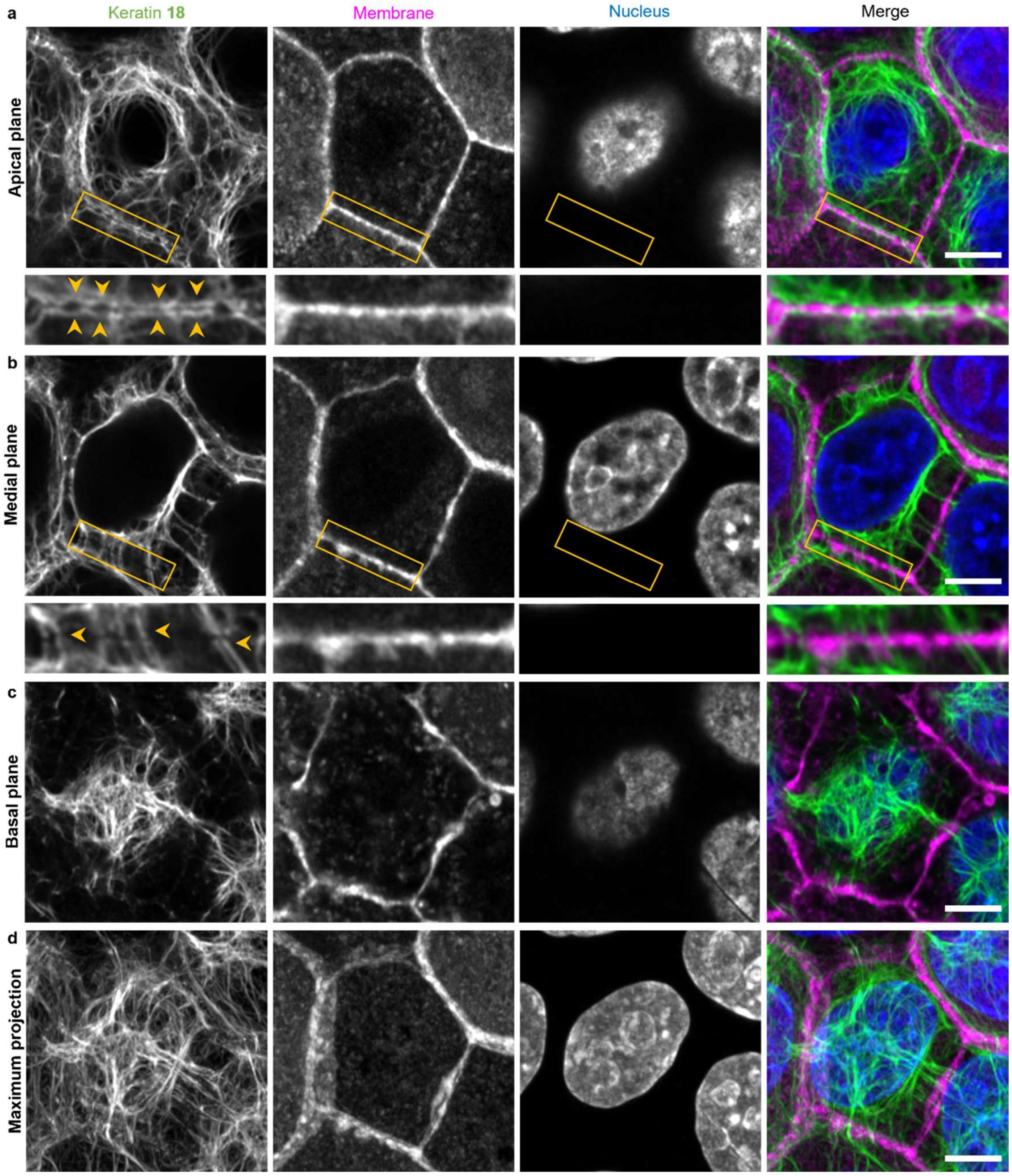


**Supplementary Figure 2: Rim-and-spoke organization in relaxed monolayers.** Fixed MDCK monolayer expressing keratin-18-GFP (green) and CAAX-mCherry (magenta) stained to label nuclei with Hoechst (blue). **a**, Apical plane. Inset: Zoom in on the keratin rim lining the plasma membrane on both sides of the cell-cell junction (yellow arrows). **b**, Medial plane. Inset: Zoom in on the keratin spokes aligning perpendicular to the membrane on both sides of the cell-cell junction (yellow arrows). **c**, Basal plane. **d**, Maximum projection of the whole cell showing that the nucleus is fully surrounded by a keratin mesh. Scale bar 5 µm.

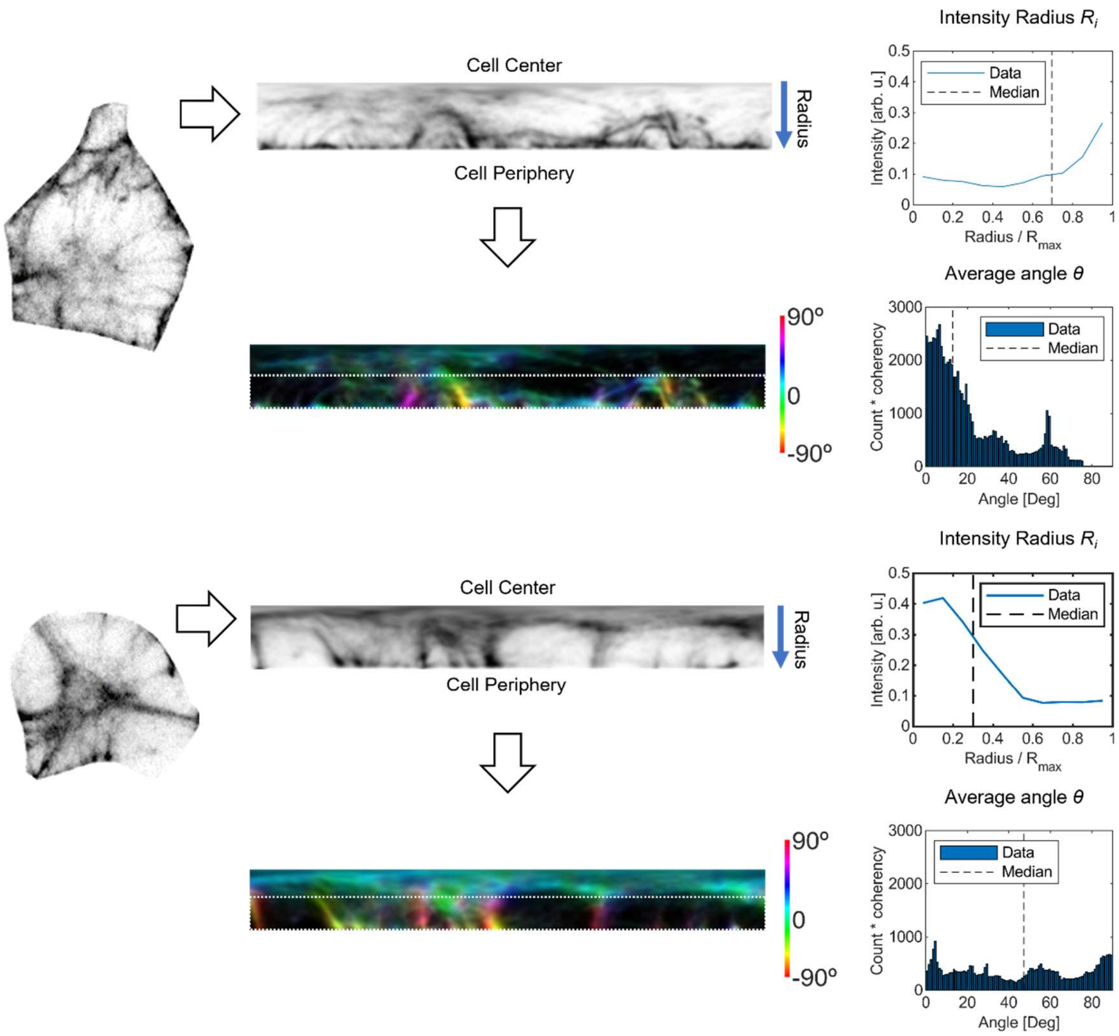


**Supplementary Figure 3: Calculation of *Ri* and *θ*.** First, the keratin signal of the cell is transformed into polar coordinates. Second, the mean intensity is measured as a function of the radius, rescaled by the maximum radius $R_{max}$. The median of this curve is $R_i$. Third, the outer half (dotted area) of the cell is used to calculate the direction of keratin features in every pixel. From this, the distributions of angles weighted by their respective coherency is determined. The median of this curve is the average angle $\theta$. For more details see the method section.

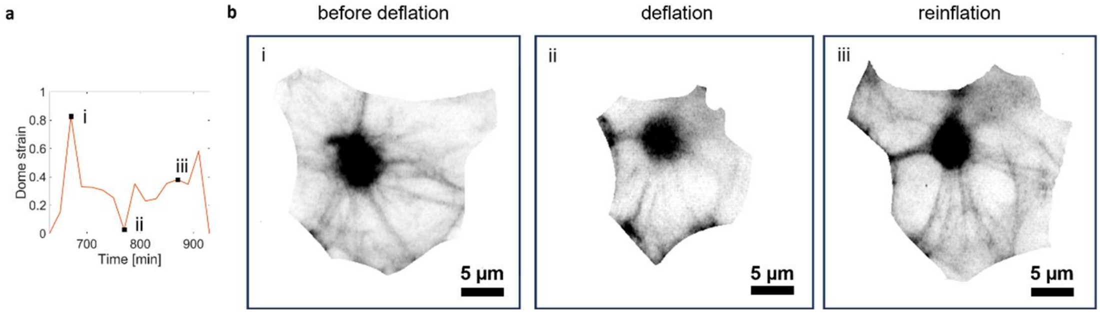


**Supplementary Figure 4: Bundles persist during deflation and reinflation. a,** Dome strain of a directed dome during a spontaneous deflation and reinflation cycle. Black squares mark the time points displayed in (**b**). **b**, Maximum projection of an individual cell expressing keratin-18-GFP showing that the star-like bundle structure persists during dome deflation and reduction in cell area.

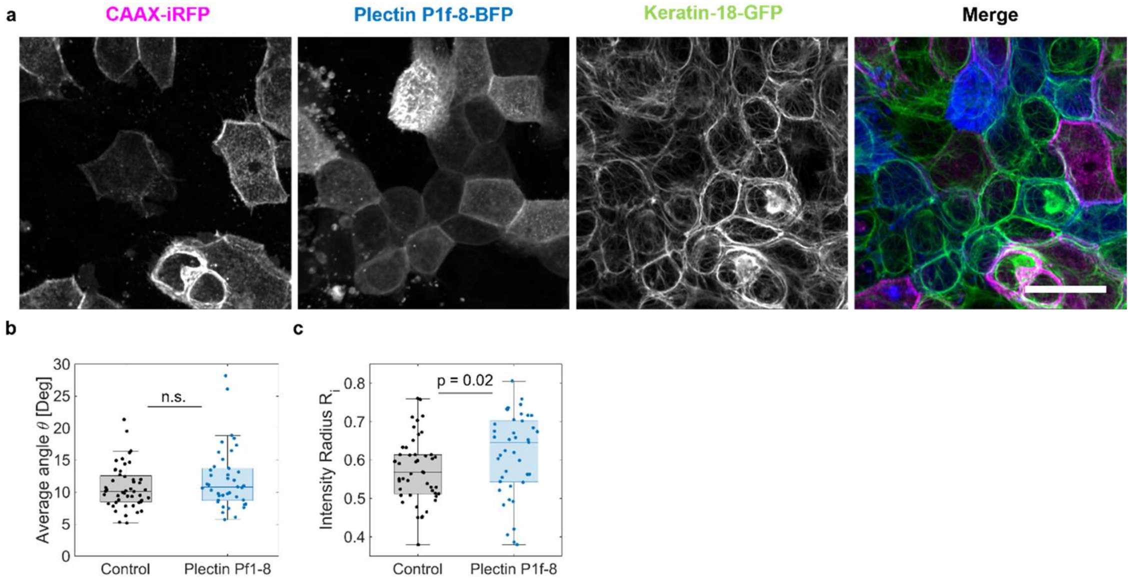


**Supplementary Figure 5: Plectin P1f-8-BPF cells display a similar keratin structure as control cells. a,** Fixed image of a mixed population of keratin-18-GFP (green) cells expressing either CAAX-iRFP (magenta, control) or P1f-8-BFP (blue). **b**, Average angle $\theta$ of control cells and cells expressing P1f-8-BFP. **c**, Intensity radius $R_i$ of control cells and cells expressing P1f-8-BFP. Measurements in (**b,c**) are based on clusters of the same cell types in mixed populations in fixed simples. Boxes indicate the median with lower and upper quartile and whiskers extend to minimal and maximal values. (n = 48 control cells, n = 40 P1f-8-BFP cells). Differences in $\theta$ are non-significant whereas differences in $R_i$ are statistically significant but small. Scale bar 20 μm.

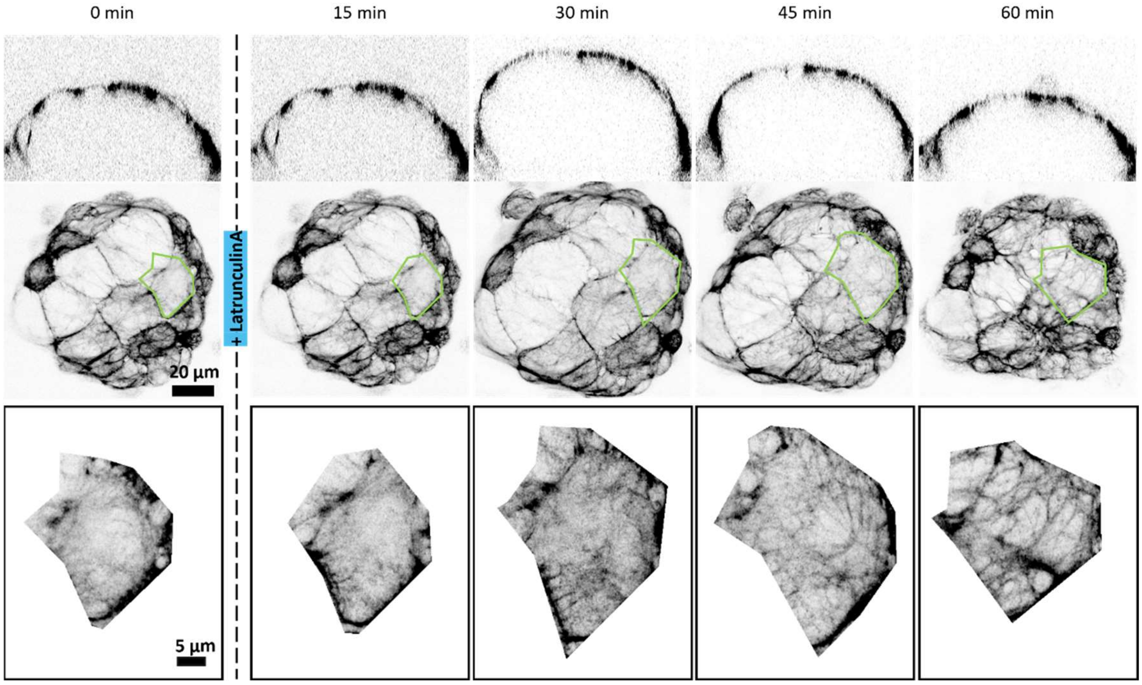


**Supplementary Figure 6: Latrunculin A treatment causes fast keratin bundling.** Top**:** Side view of an inflated dome treated with Latrunculin A. Middle: Maximum projection of the upper part of the dome with one cell segmented in green. Bottom: Individual segmented cell from the same dome. Representative images for 6 domes.

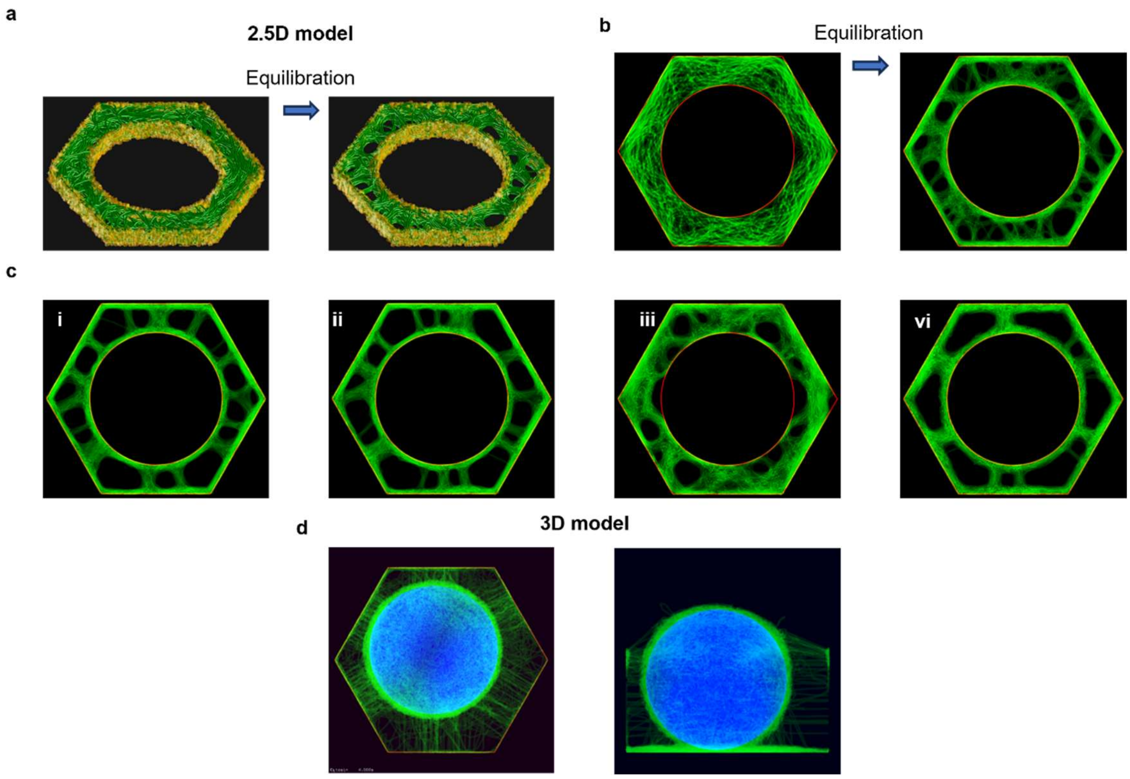


**Supplementary Figure 7: Rim-and-spoke arrangement in simulations. a,** Model view before and after equilibration showing the emergence of rim-and-spoke structure as the result of keratin (green) interaction with cortical actin and nucleus via discrete linkers (yellow) (parameters:$K^{-}/K^{+} = 0.1$, $d_b/a_0 = 0.1$). **b**, Top view of the model before and after equilibration. **c**, Alternative parameters for rim-and-spoke formation: **i)** $K^{-}/K^{+} = 1$, $d_b/a_0 = 0.1$ **ii)** $K^{-}/K^{+} = 10$, $d_b/a_0 = 0.1$ **iii)** $K^{-}/K^{+} = 0.1$, $d_b/a_0 = 0.05$ **iv)** $K^{-}/K^{+} = 10$, $d_b/a_0 = 0.2$. **d**, Top and side view of a 3D model after equilibration showing keratin (green), cell boundaries (red) and nucleus (blue).

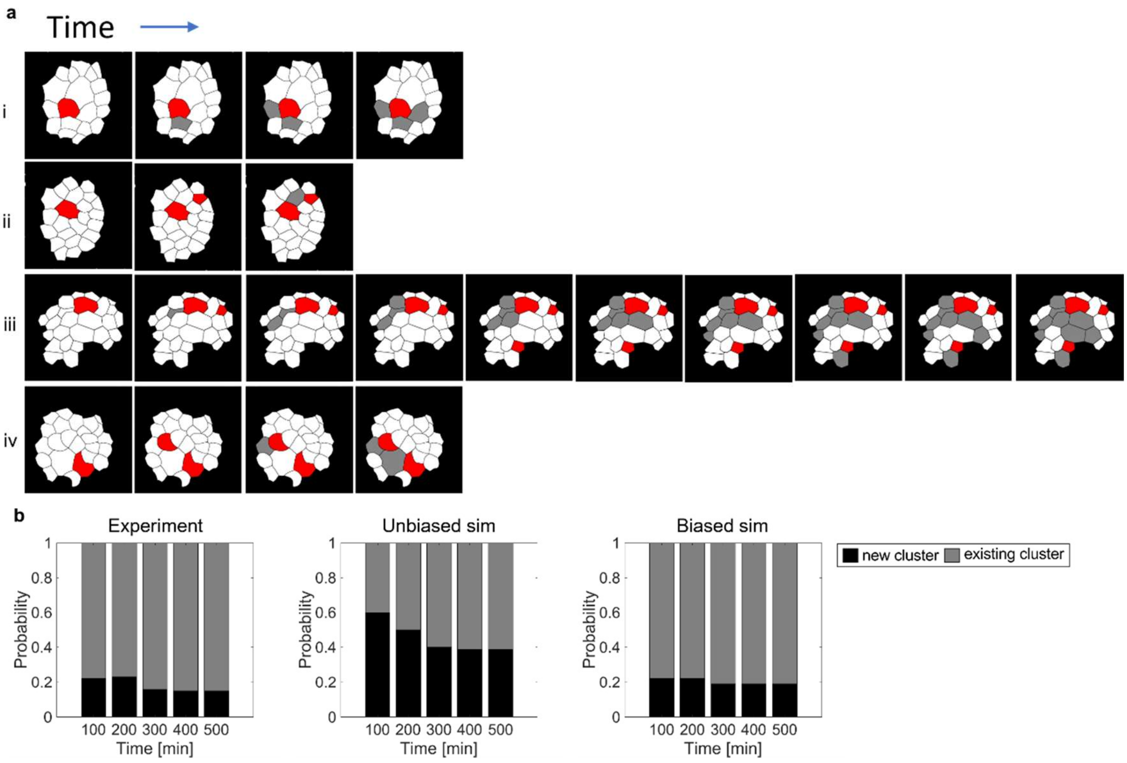


**Supplementary Figure 8: Nucleation event over time. a**, Sequence of cells crossing the bundle threshold. Cells starting a new cluster count as a nucleation event and are marked in red. Cells being connected to an existing cluster are marked in grey. Roman numbers label different experiments. **b**, Probabilities of bundling cells starting a new cluster or joining an existing cluster over time in experiments, unbiased simulations and biased simulations. The time is in reference to the first cell crossing the bundling threshold.

#### Supplementary Videos

**Supplementary Video 1: Cell in a directed dome undergoing the bundle transition.** Keratin-18-GFP signal with the result of the membrane segmentation in green.

**Supplementary Video 2: Growth of a keratin depleted zone.** Keratin-18-GFP signal with the result of the membrane segmentation in green. The field of view is a zoom on one tricellular junction of the cell displayed in supplementary video 1.

**Supplementary Video 3: Isometric view of a 3D model simulation.** A single cell with the keratin fibres in green, the nucleus in cyan and the cell borders in grey. The cell is stretched gradually up to a maximum of 1000%. During the stretch, the keratin fibres reorganize into a star-like bundle structure while the nucleus is pushed out of its keratin cage.

**Supplementary Video 4: Collective bundling process.** Cropped images of keratin-18-GFP in a directed dome. The outlines indicate segmented cell borders colour-coded according to the average angle $\theta$. The colour changes from blue for non-bundled cells to magenta for cells that crossed the bundling threshold.

### Supplementary Note

#### 2.5D computational models

Generation of entangled networks

To establish 2.5 computational models of intermediate filament reorganization in the region around the medial cell plane, we leveraged a previously described 2D model generation procedure to obtain 8 realizations of keratin networks with degree of entanglement $\mathcal{E} \approx 0.5$, as defined in reference [1]. Each model comprises 100 cylindrical and inextensible fibres interacting sterically through a harmonic potential of stiffness $k_s = 10$ nN/μm, each representing a keratin fibre (a collection of bundled filaments) of reference length $\ell_0 = 25$ μm, persistence length $\ell_p = 5$ μm, and diameter $\phi = 100$ nm. The base of the confining prism is a regular hexagon with reference apothem length $a_0 = 5$ μm and its height is $h_0 = 1.25$ μm. The nucleus is modeled as a cylindrical region of radius $r_n = 3.5$ μm which cannot be penetrated by keratin filaments, aligned with the prism height and centred with respect to the hexagonal base. The cytosolic space, corresponding to the cell region between the outer prism and the nucleus, is modelled as an environment of effective viscosity $\nu = 1$ pN s / μm$^2$ and all filament points are confined inside it by a harmonic potential whose stiffness is $k_c = 1$ nN/μm for points outside this space region and 0 for points located inside. Desmosomal junctions connecting intermediate filaments in neighbouring cells are modelled by ensuring that each fibre end remains attached to the cell lateral faces without being able to cross tricellular junctions. Following generation of the entangled networks, long-term equilibration ($t_{eq} = 100$ s) at room temperature ensures elimination of any pretension.

Formation of rim-and-spoke structures

To study the emergence of rim-and-spoke structures from initially random networks, we further allowed fibers to interact electrostatically over a distance $d_a = 100$ nm by activating an attractive potential of stiffness $k_a = 10$ pN/μm and modelled the presence of plectins on the nuclear surface and on the cell cortex. These were represented as uniformly distributed springs with a surface density of 100 μm$^{-2}$ and stiffness $k_p = 10$ pN/μm. In our models, plectins may stochastically form bonds with any fiber point located within a distance $d_b$ according to a binding rate $K^+ = 1$ s$^{-1}$ and unbind with rate $K^-$. We then let the system equilibrate for $t_{eq} = 100$ s $\gg 1/K^+$ to track the self-organization of intermediate filaments under various choices for $d_b$ (range: 0.25–1 μm) and $K^-$ (range: 0.1–10 s$^{-1}$).

Stretching of 2.5D cell models

In the steps described above, axial tension in fibres and forces on desmosomes are relatively small. Because this may not be the case during cell stretching, we enabled fibre extensibility with axial stiffness $k = 1$ nN / μm, and increased the stiffness of the confinement potential by one order of magnitude ($k_c{}' = 10$ nN/μm) to preserve the larger stiffness of desmosomal junction with respect to intermediate filaments. We also modelled plectin bonds as slip bonds, with a force-dependent unbinding rate $K_{eff}^{-} = K^{-} \exp(f/f_{ub})$, where $f_{ub} = 100$ pN is a force-sensitivity parameter. Then, we rapidly increased the cell base area by 4-fold over a time span of 0.3 s, after which the stretched configuration was maintained for 150 s. The cell height was kept constant throughout these simulations.

#### 3D computational models

Generation of entangled networks

To better investigate the interaction between keratin filaments and the nucleus, we obtained 7 realizations of fully 3D computational models comprising entangled networks ($\mathcal{E} \approx 0.5$) of 200 keratin fibres with reference length $\ell_0^{\mathrm{3D}} = 35$ μm. The nucleus is represented as a rigid sphere of radius $r_n = 3.5$ μm interacting sterically with the fibres and the cell space is a hexagonal prism of reference height $h_0^{\mathrm{3D}} = 10$ μm. All other physical parameters have the values already indicated for the generation of 2.5D models and a long-term equilibration ($t_{eq} = 100$ s) phase at room temperature is also applied to ensure elimination of any pretension within the fibre network.

Formation of rim-and-spoke structures and apical-basal asymmetry

Contrary to the 2.5D models, and motivated by the need to maintain a manageable computational cost for the 3D simulations, we represented the interaction of the fibres with the cell cortex and the nucleus by introducing a short-ranged attractive potential on both surfaces. This attracts any fibre point located inside the cell but outside the nucleus that is within a distance $d_b^{\mathrm{3D}} = 0.5$ μm from the nuclear surface with a stiffness $k_c^{\mathrm{3D}} = 10$ pN/μm. Consistently with the 2.5D simulations, we also let these models equilibrate for $t_{eq} = 100$ s to give keratin fibres time to self-organize into rim-and-spoke structures.

Then, we induced the apical-basal asymmetry that is typically observed in epithelial cells by deactivating the attractive potential acting on the top surface of the cell space enclosing the intermediate filament network and gradually halving the height of the lateral cortex over a time period of 10 s. The obtained model was then equilibrated for $t_{eq} = 100$ s.

Stretching of 3D cell models

To simulate cell loading, we enabled fibre extensibility ($k^{3D} = 10$ nN / μm), increased the stiffness of the confinement potential by one order of magnitude (${k_c}' = 10$ nN/μm), and we gradually increased the cell area to reach 1000% strain in 10 s.

#### Computational model visualization as image projections

To quantitatively analyse our computational predictions of keratin spatial distribution and compare them with experimental measurements, we used our customized version of Cytosim to extract 75-nm-thick slices of the model at each simulation time step. By treating the intermediate filament, cell cortex, and nucleus signal channels separately, we first obtained image stacks along the top-bottom (2.5D and 3D models) and front-back (3D model only) directions and then combined using Fiji's image stack projection utility[2]. "Sum slices" was used for each channel in the 2.5D models, while a maximum intensity projection was preferred for the nucleus signal in the 3D models. Regardless of the chosen method, the obtained projections were merged into composite images. These were used for model visualization and, in the case of 2.5D simulations, served as input for further quantitative analysis using the Matlab algorithm described in the "Image analysis" section.

#### Supplementary References


1. Pensalfini, M., Golde, T., Trepat, X. & Arroyo, M. Nonaffine Mechanics of Entangled Networks Inspired by Intermediate Filaments. *Phys. Rev. Lett.* **131**, 058101 (2023).
2. Schindelin, J. *et al.* Fiji: an open-source platform for biological-image analysis. *Nat. Methods* **9**, 676–682 (2012).
